# Acute hypertension causes focal coronary artery perivascular cGAS-, IRF3, and IFNAR-dependent type I interferon responses

**DOI:** 10.64898/2026.11.12.764974

**Authors:** Gilda Castellanos von Borstel, Samuel J. Wright, Betul B. Gunes, Van K. Ninh, Bo Zhang, Jessica M. Hong, Raimon Padrós-Valls, Eugene Golts, Jingjing Hu, Zhenxing Fu, Kevin R. King

**Author notes:** Corresponding author and contact: **Address for correspondence** Kevin R. King, MD, PhD, University of California San Diego, 9500 Gilman Dr. MC 0412, La Jolla, CA 92093. **Funding:** This work was funded by T32HL007444 (V.K.N; J.M.H), NIH DP2AR075321 (K.R.K), R01HL181539 (K.R.K).

## Abstract

Hypertension exposes coronary arteries to dynamic mechanical forces that alter the structural integrity and lead to pathologic changes such as aneurysm, dissection, or atherogenesis. Although the effects of chronic hypertension have been extensively studied, the early molecular responses to increased luminal pressure remain poorly understood. Here, we find that acute hypertension in murine coronary arteries induced by transverse aortic constriction (TAC) causes focal perivascular type I interferon (IFN) responses in the vessel wall in contrast to comparatively homogeneous perivascular fibrosis. Spatial transcriptomics and RNAscope independently demonstrate spatially clustered expression of interferon-stimulated genes (ISGs) in the walls of pressurized left and right coronary arteries in the first week after TAC. Genetic disruption of cGAS or IRF3 each markedly attenuated the perivascular ISG response, indicating the response is a consequence of DNA sensing. Pharmacologic disruption of type I IFN signaling using an anti-IFNAR antibody abolished pressure-induced induction of focal ISG expression suggesting the pathway can be inhibited therapeutically. Since focal type I IFN signaling in the infarct borderzone of the left ventricle increased risk of dilation and rupture, we propose that focal perivascular type I IFN signaling in the vessel wall may serve as a nidus for analogous pathologies like aneurysm, dissection, or atherogenesis and represent a potential target for therapy.

## Introduction

Coronary arteries exist within a mechanically dynamic environment in which endothelial cells, vascular smooth muscle cells, perivascular stromal cells and surrounding cardiomyocytes experience changing intraluminal pressure, shear stress, and external vascular deformation resulting from the force of myocardial contraction (PMID:34255994) (PMID:18541735). Hypertension is a known risk factor for aneurysm, dissection, arteriosclerosis, and atherogenesis (PMID:33245319) (PMID:24370755) (PMID:29472380) (PMID:136891). Elevated intraluminal pressure increases circumferential wall stress, promoting adaptive changes in vascular smooth muscle cells, extracellular matrix organization, and arterial stiffness (PMID:33245319) (PMID: 25355505) (PMID: 27118293). Although these responses may initially preserve vascular integrity, excessive or sustained mechanical loading can contribute to vascular dysfunction and susceptibility to injury (PMID:34255994). While the structural consequences of chronic hypertension have been extensively characterized (PMID: 25767286) (PMID: 33245319), the early molecular responses of coronary vessels to acute increases in intraluminal pressure remain poorly understood. In particular, whether acute mechanical stress induces localized inflammatory responses within the coronary vessel wall and surrounding perivascular tissue remains unclear.

Under conditions of excessive mechanical loading, vascular adaptive mechanisms may be disrupted, contributing to cellular injury, inflammatory activation, and extracellular matrix reorganization (PMID: 39931738) (PMID:25355505) (PMID:24663091). However, the biological consequences of increased vascular pressure may not be uniformly distributed throughout the vessel wall. Differences in local tissue architecture, cellular composition, and mechanical deformation may therefore generate spatially heterogeneous responses to increased vascular loading (PMID:18541735) (PMID:25355505). Such heterogeneity could produce focal regions of cellular stress and inflammation, potentially contributing to localized vascular vulnerability. Despite growing recognition that mechanical forces influence vascular remodeling, how acute increases in vascular pressure shape the spatial distribution of inflammatory and molecular responses within the coronary perivascular microenvironment remains poorly understood.

Understanding these localized responses requires approaches capable of resolving molecular changes within their native vascular architecture. Conventional bulk transcriptomic analyses may obscure focal molecular responses, while dissociative single-cell approaches do not preserve the anatomical relationships between vascular cells and surrounding tissues (PMID:27365449) (PMID:35859028). Spatial transcriptomics provides an opportunity to examine how transcriptional programs are organized relative to coronary vessels and whether responses to increased mechanical loading are distributed uniformly or restricted to specific vascular regions (PMID:31501547). In particular, examining the radial distribution of molecular programs relative to the vessel wall and their angular organization around the vessel circumference may reveal localized cellular responses that would otherwise remain undetected (PMID:27365449). Defining these spatial relationships may provide new insight into how coronary vascular stress is translated into heterogeneous molecular responses within the surrounding perivascular microenvironment.

Here, we combined transverse aortic constriction (TAC), Xenium spatial transcriptomics, multiplexed RNA in situ hybridization, and genetic and pharmacological perturbations to investigate the early structural and molecular responses of coronary vessels to increased mechanical loading. We found that pressure overload in the coronary arteries not only induces perivascular collagen accumulation but also an unexpected focal and circumferentially asymmetric type I interferon (IFN) response characterized by expression of interferon-stimulated genes (ISGs). Genetic disruption of cyclic GMP-AMP synthase (cGAS) or interferon regulatory factor 3 (IRF3) markedly attenuated perivascular induction of the ISG, *Ifit1*, while pharmacological blockade of the type I interferon receptor (IFNAR) strongly suppressed this response. Despite inhibition of IFNAR-dependent ISG expression, ISG suppression was not accompanied by a detectable reduction in early perivascular collagen deposition under the conditions tested. Together, these findings identify a focal, pharmacologically modifiable innate immune response associated with acute coronary pressure overload. We propose that focal activation of this pathway may represent an early molecular response to vascular mechanical stress, with potential implications for understanding the processes that precede overt arterial injury and structural failure.

## Results

### Hypertensive stress induces bilateral coronary artery perivascular fibrosis

To determine whether pressure overload alters the coronary perivascular environment, we performed transverse aortic constriction (TAC) in 12-week-old wild-type C57BL/6J mice and analyzed hearts at day 7 (Fig. 1A). TAC heart samples were accompanied by an increase in the cardiac stress-associated transcript *Nppa* visualized by duplex RNA *in situ* hybridization and quantified as *Nppa*-positive tissue area (Fig. 1B). Histological analysis revealed increased collagen accumulation following TAC (Fig. 1C). Quantification showed significant increase in left ventricular interstitial fibrosis, whereas right ventricular interstitial fibrosis did not differ significantly between controls and the measured endpoints (Fig. 1C-D; Supplementary Fig. 1). These findings support a stress-associated response from TAC to the left ventricle. High-magnification imaging of vessels in left and right coronary arteries (LCA and RCA) revealed increased perivascular collagen deposition (Fig. 1E-F), confirmed by perivascular area-based quantification in both LCA and RCA (Fig. 1F). Together, these results establish that mechanical stress imposed by TAC produces bilateral coronary arterial perivascular fibrosis.

**Figure 1.** Hypertensive stress induces bilateral coronary artery perivascular fibrosis. **(A)** Schematic illustration of transverse aortic constriction (TAC) model used to induce mechanical stress on the myocardium and coronary vasculature through pressure overload. TAC increases proximal aortic and coronary arterial pressure, resulting in increased luminal stress on the coronary vessel wall. **(B)** Representative whole-heart RNAscope images showing *Nppa* expression in control and wild-type (WT) TAC hearts, with corresponding quantifications of *Nppa*-positive area expressed as percentage of total tissue area. Scale bar, 1 mm. **(C)** Whole-heart Masson’s trichrome-stained sections from control and WT TAC hearts illustrating increased myocardial collagen deposition (blue) after imposed stress. Scale bar, 1 mm. **(D)** Quantification of interstitial collagen-positive area in the left ventricle (LV) and right ventricle (RV), expressed as a percentage of myocardial area. **(E)** Representative high-magnification Masson’s trichrome-stained images of left coronary artery (LCA) and right coronary artery (RCA) from control and WT TAC hearts. Scale bars, 50 μm. **(F)** Quantification of perivascular collagen-positive area surrounding the LCA and RCA, expressed as a percentage of the perivascular area. Bars represent mean ± SEM, with individual data points representing individual arteries. Statistical comparisons were performed using unpaired, two-tailed Student’s t-test or one-way ANOVA, as appropriate. Statistical significance is indicated as **P < 0.001, ***P < 0.0001, and ns as non-significant. Exact sample sizes and statistical details are provided in the corresponding Methods section.

### Vessel-centered spatial transcriptomics reveals focal interferon-stimulated gene niches surrounding coronary arteries

We next asked what molecular programs are activated within the wall of mechanically stretched coronary arteries. We collected hearts on day 7 or day 28 after TAC or from non-TAC controls and performed Xenium spatial transcriptomic profiling using a 5,050-probe panel (n = 2 control, n = 2 TAC day 7 and n = 3 TAC day 28). Whole-heart transcript mapping revealed the anatomical distribution of cardiomyocyte-, fibroblast-, and smooth muscle-associated transcripts, enabling the identification of coronary vascular structures. Using a smooth muscle transcript-guided vessel detection and hull-tracing workflow, we identified 60 intramyocardial vessels across the dataset (Fig. 2A). Visualization of smooth muscle-associated transcripts across all identified vessels demonstrated the range of vascular structures included in the analysis, comprising 13 control and 47 TAC vessels (Supplementary Fig. 2).

**Figure 2.** Vessel-centered spatial transcriptomics reveals focal ISG niches surrounding coronary arteries. **(A)** Schematic representation of the computational approach used to identify and segment coronary vessels in spatial transcriptomic datasets. Coronary vessels were identified using vascular smooth muscle-associated transcripts, followed by segmentation of the vessel lumen and estimation of the surrounding perivascular boundary. **(B)** Schematic illustrating vessel-centered radial and angular analyses. Radial analysis quantifies gene expression as a function of distance from the vessel center using concentric 100-μm intervals, while angular analysis evaluates the circumferential distribution of gene expression relative to the vessel center. **(C)** Representative radial distributions of individual genes (left) and transcriptional programs identified by Gaussian mixture modeling (GMM; right). Smooth muscle-associated genes (*Myh11, Tagln*) and the corresponding smooth muscle transcriptional program are concentrated near the vessel center, whereas fibroblast-associated genes (*Col1a2, Lum*) and cardiomyocyte-associated genes (*Myh6, Myl2*) exhibit broader radial distributions. Radial transcriptional programs include smooth muscle (SMC), fibrosis-associated, and ventricular cardiomyocyte-associated programs (vCM). **(D)** Representative spatial maps showing the anatomical localization of vascular smooth muscle (*Myh11, Tagln*), perivascular fibroblast (*Col1a2, Lum*), and ventricular cardiomyocyte (*Myh6, Myl2*) transcriptional signatures relative to the coronary vessel (H&E). **(E)** Quantification of program-associated area occupied by proliferation-, macrophage-, and fibroblast-associated transcriptional programs in control and TAC arteries. P values were calculated using two-sided Mann–Whitney U tests. **(F)** Genome-wide ranking of 3,335 genes according to angular concentration, quantified using mean resultant length (R). Higher R values indicate greater angular concentration, whereas lower values indicate more diffuse circumferential distributions. Interferon-stimulated genes, including *Ifit1, Ifit3, Isg15, Cxcl10, and Usp18*, are among genes exhibiting elevated angular concentration. **(G)** Representative angular distributions of transcriptional programs, including smooth muscle, fibrosis-associated, ventricular cardiomyocyte-associated, and ISG-associated programs. **(H)** Angular distributions of individual interferon-stimulated genes *(Ifit1, Ifit3, Isg15, Cxcl10, and Usp18)* within a representative coronary vessel. Spatial transcriptomic analysis included 60 intramyocardial vessels identified across seven mice (control, n = 2; TAC day 7, n = 2; TAC day 28, n = 3), comprising 13 control and 47 TAC vessels.

We next developed a vessel-centered framework to distinguish radial transcriptional programs organized by distance from the arterial vessel wall from angular programs enriched within discrete circumferential sectors (Fig. 2B). To account for noncircular vessel geometry, vessel wall boundaries were estimated from smooth muscle-associated transcripts in successive 5° angular sectors, allowing transcript distances to be normalized to the local vessel wall radius (Supplementary Fig. 3). Radial modeling recovered expected smooth muscle (Myh11, Tagln), fibroblast/extracellular matrix (Col1a2, Lum), and cardiomyocyte (Myh6, Myl2)-associated transcripts (Fig. 2C). These distributions corresponded to the expected anatomical organization of vascular smooth muscle, surrounding fibroblasts, and adjacent myocardium (Fig. 2D). To identify transcriptional programs shared across arteries, we compared vessel-specific gene clusters using a network-based approach and Louvain community detection, identifying 16 transcriptional metaclusters (Supplementary Fig. 4). Quantification of the spatial area occupied by each transcriptional program revealed greater coverage of proliferation-, macrophage-, and fibroblast-associated programs in TAC vessels compared with controls (Fig. 2E; Supplementary Fig. 5). Together, these analyses demonstrate that the vessel-centered approach captures established vascular organization and spatially heterogeneous transcriptional responses within the coronary microenvironment.

We next examined whether transcriptional programs exhibited preferential localization around the vessel circumference. Angular analysis identified an interferon-stimulated gene (ISG) program characterized by I*fit1, Ifit3, Isg15, Cxcl10, and Usp18* (Fig. 2F). Unlike the broader angular distributions of smooth muscle-, fibroblast-, and cardiomyocyte-associated programs, the ISG program exhibited a pronounced angular concentration (Fig. 2G). Radial analysis of individual ISGs further demonstrated enrichment at defined distances from the vessel center (Fig. 2H). Together, these findings identify an unexpected focal ISG-associated transcriptional program within the coronary perivascular microenvironment following TAC.

Having identified a focal ISG-associated transcriptional program through vessel-centered analysis, we next examined its anatomical distribution within the coronary vasculature. Whole-heart Xenium imaging demonstrated the distribution of *Ifit1* and *Ifit3* transcripts across the myocardium and their localization relative to smooth muscle-associated *Myh11* expression (Fig. 3A). Higher-magnification analysis of coronary arteries from TAC hearts revealed focal enrichment of *Ifit1* and *Ifit3* transcripts within the perivascular microenvironment, with an asymmetric distribution around the vessel circumference (Fig. 3B). We also examined human cardiac Xenium data and observed localized expression of *IFIT1* and *IFIT3* transcripts within a vascular region, including their distribution relative to *MYH11*-expressing smooth muscle cells (Fig. 3C). Identification of focal perivascular ISG expression in both mouse and human cardiac tissue suggests that this localized interferon-associated response represents a shared response by mechanically stressed cardiac vasculature.

**Figure 3.** Spatially localized ISG expression in mouse and human cardiac vessels. **(A)** Representative whole-heart mouse Xenium spatial transcriptomic images showing H&E staining (left), Ifit1 and Ifit3 transcripts (magenta; middle), and their distribution relative to the vascular smooth muscle-associated transcript *Myh11* (purple; right). Dashed boxes in the H&E image indicate the left coronary artery (LCA) and right coronary artery (RCA) regions. LV, left ventricle; RV, right ventricle. Scale bars, 5,000 μm. **(B**) Representative mouse images of a coronary artery 7 days post TAC. H&E staining (left) illustrates vessel morphology, while spatial transcript (Xenium) maps show the distribution of ISG-associated transcripts (*Ifit1* and *Ifit3*; magenta) alone (middle) or together with the vascular smooth muscle-associated transcript *Myh11* (purple; right). Dashed outlines indicate the vessel lumen. Scale bars, 200 μm. **(C)** Representative human Xenium spatial transcriptomic images showing H&E staining (left), ISG-associated transcripts *IFIT1* and *IFIT3* (magenta; middle), and their distribution relative to the smooth muscle-associated transcript *MYH11* (purple; right). Dashed outlines indicate the vessel lumen. Scale bars, 200 μm. **(D)** Representative RNAscope images showing *Ifit1* expression in the left coronary artery (LCA; top) and right coronary artery (RCA; bottom) territories of control and wild-type TAC mice. Scale bars, 50 μm. **(E)** Quantification of Ifit1-positive area normalized to total perivascular area in the LCA (top) and RCA (bottom) of control and TAC mice. Data are presented as mean ± SEM, with individual points representing biological replicates. Statistical significance is indicated as ***P < 0.001 and ****P < 0.0001. Exact sample sizes and statistical details are provided in the corresponding Methods section.

We next used RNAscope to independently validate the induction of perivascular ISGs following TAC. Control hearts exhibited minimal *Ifit1* signal, whereas TAC hearts displayed prominent focal *Ifit1* expression surrounding coronary vessels (Fig. 3D). Quantification demonstrated significantly increased *Ifit1*-positive area within the perivascular regions of both left and right coronary artery territories (Fig. 3E). Together, these findings demonstrate that TAC induces focal perivascular *Ifit1* expression in both coronary territories, consistent with the ISG program identified by spatial transcriptomics.

### cGAS-IRF3 regulate the coronary perivascular ISG niches following mechanical stress

Having identified a focal perivascular ISG expression following mechanical stress, we next investigated the molecular pathway responsible for its induction. Cytosolic DNA sensing by cyclic GMP–AMP synthase (cGAS) can activate STING-dependent signaling and the transcription factor IRF3, promoting type I interferon production (PMID:23258413) (PMID:23258412) (PMID:19776740); secreted IFNs can then stimulate neighboring cells to express interferon-stimulated genes via their cognate cell surface receptor IFNAR (PMID:24555472) (PMID:24766893). Previous work in mechanically stressed myocardial injury in the infarct borderzone or at sites of sterile needle injury implicated cGAS-dependent interferon signaling in the heart. We hypothesized that the acute luminal pressure induced by TAC caused distension and mechanical stress resulting in focal microinjuries within the wall resulting in an analogous activation of the cGAS–STING–IRF3 signaling axis, leading to type I IFN production and subsequent IFNAR-dependent upregulated expression of ISG effector molecules (Supplementary Fig. 7) (PMID:27013428) (PMID:25023467). To test this hypothesis, we examined the perivascular ISG response following TAC in mice deficient in either cGAS or IRF3. Whole-heart duplex RNAscope demonstrated widespread *Nppa* expression in TAC hearts across all genotypes, whereas perivascular Ifit1 expression was markedly attenuated in IRF3-deficient (*Irf3*^−/−^) and cGAS-deficient (*Cgas*^−/−^) mice compared with wild-type (WT) TAC controls (Fig. 4A). Higher-magnification imaging further demonstrated reduced focal Ifit1 expression within the perivascular regions of both left and right coronary artery territories in both knockout groups (Fig. 4B–D). In contrast, perivascular *Nppa* expression did not differ significantly between genotypes, suggesting that attenuation of the ISG response occurred despite a persistent myocardial stress-associated transcriptional response (Supplementary Fig. 8). Together, these loss-of-function experiments demonstrate that cGAS and IRF3 are required for robust perivascular *Ifit1* induction following coronary artery hypertension.

**Figure 4.** cGAS and IRF3 are required for robust coronary perivascular Ifit1 induction following mechanical stress. **(A)** Representative whole-heart RNAScope images showing *Nppa* (cyan) and *Ifit1* (magenta) expression in WT, Irf3 knockout (*Irf3*⁻/⁻), and Cgas knockout (*Cgas*⁻/⁻) hearts following TAC. *Nppa* expression is observed across all three genotypes, consistent with a cardiac stress response. Scale bars, 1 mm. **(B)** Representative high-magnification RNAscope images showing *Nppa* (cyan) and *Ifit1* (magenta) expression surrounding coronary vessels in the left coronary artery (LCA) and right coronary artery (RCA) territories in WT, *Irf3*⁻/⁻, and *Cgas*⁻/⁻ TAC hearts. Scale bars, 50 μm. **(C–D)** Quantification of Ifit1-positive area surrounding the LCA **(C)** and RCA **(D),** expressed as a percentage of the perivascular area. Perivascular Ifit1 expression is markedly reduced in Irf3^-/-^ and Cgas^-/-^ mice compared with WT mice following TAC. Data are presented as mean ± SEM, with individual data points representing individual vessels. Statistical analyses were performed using one-way ANOVA, ****P < 0.0001. Exact sample sizes and statistical details are provided in the corresponding Methods section.

To test whether cGAS and IRF3 loss alters early perivascular collagen deposition, we assessed collagen deposition using Masson’s trichrome staining in WT, *Irf3*^−/−^, and *Cgas*^−/−^ hearts following TAC. Whole-heart and high-magnification images revealed perivascular collagen accumulation across all three genotypes (Supplementary Fig.9A-B). Quantification demonstrated no significant differences in perivascular fibrosis between WT TAC and either knockout group in the LCA or RCA territories (Supplementary Fig. 9C–D). Thus, despite markedly attenuating perivascular *Ifit1* induction, genetic disruption of cGAS or IRF3 did not significantly alter early perivascular collagen accumulation. These findings suggest that cGAS- and IRF3-dependent ISG activation are not required for early perivascular fibrotic deposition following mechanical stress.

### IFNAR signaling is required for coronary perivascular ISG induction following mechanical stress

To evaluate the therapeutic potential of inhibiting the hypertension-induced perivascular IFN response, we tested whether ISG expression could be pharmacologically inhibited at the time of acute hypertensive stress. To determine whether the observed ISG response required type I IFN receptor signaling and the IFN alpha receptor (IFNAR), we pharmacologically blocked it 2 days prior to TAC using an established mouse anti-IFNAR neutralizing antibody (Fig. 5a) (PMID:39198639) (PMID:29106401), analogous to a human biologic FDA approved for lupus (PMID: 31851795). RNAscope demonstrated marked reduction of perivascular Ifit1 expression following anti-IFNAR treatment compared with isotype-treated controls (Fig. 5B). Quantification confirmed significantly decreased *Ifit1*-positive perivascular area in both left and right coronary artery territories following IFNAR blockade (Fig. 5C). In contrast, *Nppa* expression did not differ significantly between treatment groups, suggesting that suppression of the perivascular ISG response occurred without a detectable reduction in the measured myocardial stress-associated transcriptional response (Supplementary Fig. 10). These findings demonstrate that perivascular Ifit1 induction depends on IFNAR signaling and can be pharmacologically suppressed, identifying a potentially targetable component of the vascular response to pressure overload.

**Figure 5.** IFNAR signaling is required for coronary perivascular ISG induction following mechanical stress. **(A)** Schematic of experimental design. Mice received isotype control or anti-IFNAR antibody (500 μg, intraperitoneally) on days −2, +2, +4, and +6 relative to transverse aortic constriction (TAC), with tissue collected on day 7. (B) Representative RNAscope images showing *Nppa* (cyan) and *Ifit1* (magenta) expression in the left coronary artery (LCA) and right coronary artery (RCA) territories of TAC mice treated with isotype control or anti-IFNAR antibody. Scale bars, 50 μm. **(C)** Quantification of *Ifit1*-positive area expressed as a percentage of the perivascular area in LCA and RCA territories of isotype control or anti-IFNAR antibody treated mice. **(D)** Quantification of *Nppa*-positive area expressed as a percentage of the perivascular area in LCA and RCA territories of isotype control or anti-IFNAR antibody treated mice. Each dot represents individual vessel. Data are presented as mean ± SEM. Statistical significance was determined using one-way ANOVA. ns, not significant; ****P < 0.0001. Exact sample sizes and statistical details are provided in the corresponding Methods section.

Despite the marked suppression of perivascular *Ifit1* expression, IFNAR blockade did not significantly reduce perivascular collagen deposition surrounding either coronary territory (Supplementary Fig. 11). Together, these findings suggest that early perivascular ISG induction and collagen deposition may represent partially distinct responses to mechanical stress, with IFNAR-dependent signaling not required for the full extent of collagen accumulation observed at this time point.

## Discussion

Here, we identify a focal type I IFN response within the coronary perivascular microenvironment following acute hypertensive stress. Although hypertension imposes a broad hemodynamic load on coronary vessels, the resulting molecular responses are not uniformly distributed throughout the vascular wall. Vessel-centered spatial transcriptomics revealed focal and asymmetric enrichment of interferon-stimulated genes (ISGs) surrounding coronary arteries, while RNAscope independently confirmed increased perivascular *Ifit1* expression in both coronary territories. These findings extend previous work identifying spatially organized cGAS-IRF3-type I interferon responses after myocardial infarction (PMID:39198639) (PMID:29106401), suggesting that localized innate immune signaling may be a recurring feature of cardiovascular tissue stress. Importantly, the heterogeneous distribution of ISGs raises the possibility that increased vascular loading produces distinct local tissue responses, potentially reflecting differences in mechanical deformation, cellular injury, or the surrounding microenvironment (PMID:27013428) (PMID:39088918). Such focal inflammatory responses may identify regions of heightened vascular stress that would otherwise remain undetected using conventional approaches.

Previous studies have demonstrated that mechanical injury can activate cGAS-dependent DNA sensing and downstream type I interferon signaling (PMID:23258413) (PMID:23722159) (PMID:33927185), providing a plausible mechanism by which increased intraluminal stress could initiate localized inflammatory responses. Consistent with this model, global loss of cGAS or IRF3 markedly attenuated perivascular *Ifit1* induction following TAC, while pharmacological blockade of IFNAR strongly suppressed the response in both coronary territories. These findings support a model in which mechanical stress engages cGAS–IRF3-associated innate immune signaling and downstream IFNAR-dependent transcriptional responses, although the initiating cellular injury, DNA source, and interferon-producing cell populations remain to be established (PMID:25642965) (PMID:25023467). Perivascular fibroblasts represent a plausible initiating population given their location within the coronary perivascular microenvironment and their sensitivity to mechanical stress and changes in extracellular matrix tension. Alternatively, fibroblasts may respond to interferons produced by neighboring vascular or immune cells, amplifying the local ISG response through IFNAR signaling. Future studies using fibroblast-specific disruption of cGAS and IFNAR signaling, together with spatial assessment of nuclear injury and cytosolic DNA accumulation, will be necessary to determine whether perivascular fibroblasts initiate or propagate this response.

Our findings raise the possibility that focal perivascular interferon signaling represents an early molecular response to coronary mechanical stress, with potential implications for diseases characterized by localized arterial wall injury. Spontaneous coronary artery dissection (SCAD) is a particularly relevant clinical condition, characterized by separation of the coronary arterial wall in the setting of underlying arterial susceptibility and potential mechanical triggers (PMID:29481627; PMID:32256714). Although the mechanisms underlying coronary arterial vulnerability remain incompletely understood, our findings identify a localized innate immune response that may contribute to the vascular response to mechanical stress. We hypothesize that focal ISG induction may initially reflect vascular stress or microinjury but, if sustained or amplified, could contribute to local inflammation and progressive arterial wall vulnerability, potentially increasing susceptibility to dissection. The observation of focal vascular-associated ISG expression in human cardiac tissue suggests that similar spatially organized inflammatory responses may occur beyond experimental mouse models. Thus, whether this response contributes to arterial wall injury or precedes dissection remains unknown. Future studies examining the temporal relationship between focal ISG activation, vascular injury, and arterial wall integrity will be necessary to determine its potential role in SCAD pathogenesis.

The ability to pharmacologically suppress perivascular ISG expression through IFNAR blockade provides a potential therapeutic strategy for limiting interferon-associated inflammation in mechanically stressed coronary vessels. The clinical feasibility of targeting this pathway is supported by the use of anifrolumab, a monoclonal antibody against IFNAR1 approved for systemic lupus erythematosus, demonstrating that type I interferon signaling can be therapeutically inhibited in humans (PMID:31851795; PMID:36369793; PMID:35016780; PMID:37399470). These findings raise the possibility that IFNAR-directed therapies could be investigated to mitigate inflammatory responses in coronary vessels susceptible to mechanical injury, including those affected by SCAD. Whether suppression of this response preserves arterial wall integrity, reduces susceptibility to dissection, or limits secondary inflammatory injury remains unknown. Future studies examining the effects of IFNAR inhibition on vascular injury and repair will be necessary to determine whether targeting this pathway provides therapeutic benefit. Together, our findings identify IFNAR-dependent signaling as a pharmacologically modifiable component of the coronary response to pressure overload and a potential therapeutic target for vascular diseases characterized by focal arterial injury.

## Conclusion

Here, we identify a focal perivascular interferon-stimulated gene response to acute coronary pressure overload that depends on cGAS and IRF3 for robust induction and can be pharmacologically suppressed through IFNAR blockade. These findings reveal a previously unrecognized connection between coronary mechanical stress and localized innate immune signaling, suggesting that focal ISG activation is a candidate mechanism to regional vascular vulnerability and susceptibility to injury. In particular, this response may have implications for diseases characterized by focal coronary arterial wall disruption, including spontaneous coronary artery dissection (SCAD). Although its contribution to vascular injury remains to be established, the ability to suppress this response through IFNAR inhibition identifies a potentially actionable pathway for investigating and ultimately mitigating inflammation-associated coronary vascular injury.

## Methods

### Animals

All animal experiments were conducted in accordance with protocols approved by the University of California San Diego (UCSD) Institutional Animal Care and Use Committee (IACUC; protocol S17144). Wild-type C57BL/6J (000664), IRF3-deficient (Irf3^−/−^) were bred in house from existing colonies. cGAS-deficient (Cgas^−/−^, 026554) mice were obtained from The Jackson Laboratory and bred in-house. Experiments were performed using 12-week-old male mice, with experimental groups matched for age and sex at 12-weeks. Animals were not randomly assigned to experimental groups, and sample sizes were not determined by a priori power calculations. Scientists conducting the experimental analysis were blinded to experimental conditions

### Transverse Aortic Constriction (TAC) surgery

Transverse aortic constriction (TAC) was performed in 12-week-old mice. Mice were anesthetized and maintained under continuous ventilation. Following thoracic opening, the transverse aorta was exposed and constricted using a synthetic suture ligature tied around the aorta and a 27-gauge needle to standardize the degree of constriction. The needle was subsequently removed, leaving a calibrated aortic narrowing. Following surgery, mice were monitored for 5 days and treated with appropriate analgesic. At 7 or 28 days following TAC, hearts were collected using PBS perfusion. Cardiac tissues were fixed in 5% formalin and paraffin-embedded for subsequent histological and spatial transcriptomic analyses.

### IFNAR antibody administration

Adult C57BL/6J mice received four intraperitoneal injections of 500 μg of a neutralizing monoclonal antibody against the type I interferon receptor (IFNAR; clone MAR1-5A3; Bio X Cell, catalog no. BE04241), administered 2 days before TAC and on days 2, 4, and 6 following surgery. Control mice received an equivalent dose of isotype Ab control according to the same administration schedule. At day 7 following TAC, mice were euthanized, and hearts were collected for subsequent histological and RNAscope analyses.

### Tissue preparation and histology

Hearts were PBS perfused, fixed in 10% formalin for 12 hours, embedded in paraffin, and sectioned at 7μm thickness. Sections were collected at anatomically matched ventricular levels to permit comparisons of left and right coronary vascular territories. Collagen was visualized using Masson’s trichrome according to manufacturer’s protocol. Coronary vessels were identified based on anatomical location. Images were acquired using VS200 Scanner B under identical acquisition conditions. For each vessel, the vascular lumen and outer vascular wall were segmented. A standardized perivascular region of interest (ROI) was generated extending 100 μm from the outer vessel boundary. Perivascular fibrosis was calculated as collagen-positive area within the perivascular ROI divided by total analyzable perivascular ROI area. Regions occupied by the lumen and media were removed before calculation. Adjacent interstitial ROIs were defined using the exact rule. Image analysis was performed using QUPath and ImageJ, with blinded thresholds based on their group.

### In situ hybridization

Chromogenic duplex RNAscope *in situ* hybridization was performed on paraffin sections according to the manufacturer’s protocol. Probes targeting *Ifit1*(Cat No. 500071-C2) and *Nppa* (Cat No. 1555751-C1) were hybridized to tissue sections. Positive and negative control probes were included to assess RNA quality and nonspecific signal. Images were acquired using the VS200 Scanner B under identical settings across experimental groups. Perivascular ROIs were registered to corresponding vascular structures using anatomical landmarks. Vascular regions were matched based on vessel morphology and anatomical position. *Ifit1*- and *Nppa*-positive area were calculated as probe-positive area divided by total ROI area.

### Human tissue

Human left atrial appendage (LAA) tissue for Xenium spatial transcriptomic analysis was obtained from 14 patients, including 11 undergoing cardiac surgery and 3 undergoing lung surgery, at UC San Diego in collaboration with the Department of Pathology (Dr. Jingjing Hu). LA enlargement was assessed using clinical echocardiographic reports; controls had no echocardiographic evidence of LA enlargement. Formalin-fixed, paraffin-embedded tissue sections were used for Xenium analysis. Specimens were collected under UC San Diego IRB-approved protocol 810422, with [informed consent or an IRB-approved waiver—confirm]. Representative vascular regions shown originated from LA wall.

### Xenium Prime In Situ Gene Expression

Mouse cardiac formalin-fixed, paraffin-embedded (FFPE) tissue sections (7 µm) were processed using the Xenium Prime In Situ Gene Expression workflow (10x Genomics), following the manufacturer’s protocol (CG000580). The Xenium Prime 5K Mouse Pan Tissue and Pathways Assay Kit was supplemented with a Xenium Prime 5K Custom Add-On Panel (PN-1000731). Custom priming oligonucleotides and probes were resuspended in TE buffer (pH 8.0). Following deparaffinization and decrosslinking, slides were assembled in Xenium Cassettes v2 (PN-1000723). Target RNA was hybridized with priming oligonucleotides, followed by RNase treatment, overnight hybridization (16–24 h) with Xenium Prime DNA probes, ligation, and in situ amplification of circularized probes.

Cell segmentation staining was performed using Xenium Cell Segmentation Staining Reagents (PN-1000661). Sections were incubated overnight with Xenium Multi-Tissue Stain Mix following blocking, then treated with Xenium Stain Enhancer and Autofluorescence Mix. Nuclei were counterstained with DAPI. Slides were imaged on the Xenium Analyzer (10x Genomics) using automated cycles of fluorescent probe hybridization, imaging, and probe removal. Regions of interest were defined on the field-of-view (FOV) grid, and FOVs with less than approximately 5% tissue coverage were excluded. Barcodes were decoded using the panel design ID associated with each slide. Multimodal cell segmentation was performed using nuclear, membrane, and interior staining channels, and decoded transcripts were assigned to segmented cells.

#### Spatial transcriptomics

Spatial transcriptomic profiling was performed using the 10x Genomics Xenium platform with a 5,050-probe panel to characterize gene expression within the coronary perivascular microenvironment. Ventricular tissue sections were analyzed from seven mice, including sham-operated controls (n = 2), mice collected 7 days after transverse aortic constriction (TAC; n = 2), and mice collected 28 days after TAC (n = 3). Spatial transcript coordinates were used to identify coronary vessels based on the anatomical organization of vascular smooth muscle-associated transcripts. An image-based vessel identification workflow, implemented using the OpenCV (cv2) Python package and supplemented by manual curation, identified 60 vessels across the seven samples for subsequent vessel-centered spatial analyses. Details of vessel identification, spatial coordinate transformation, and downstream transcriptional program analysis are described below.

#### Coronary vessel identification

Coronary vessels were identified from Xenium spatial transcriptomic data using a computational workflow implemented with the OpenCV (cv2, v4.13.0.92) Python package. Ventricular regions were first delineated using the spatial distribution of cardiomyocyte-associated transcripts (*Myl2 and Myh7*), and transcripts located within these regions were retained for subsequent analysis. Vascular structures were then identified based on the spatial organization of smooth muscle-associated transcripts (*Myh11, Tagln, Cnn1, Fam129a, Col18a1*). An automated contour-detection algorithm was used to identify enclosed, approximately elliptical structures formed by these transcripts and generate boundaries corresponding to putative vascular lumens. Candidate structures with an enclosed area greater than 50 μm² were retained for further evaluation. Transcript visualization parameters and area thresholds were manually adjusted when necessary to accommodate differences in transcript density between samples. Candidate vessels were visually inspected, and structures inconsistent with coronary arterial anatomy were excluded from downstream analysis. Vessel center coordinates were subsequently extracted and transformed into the original Xenium coordinate system using coordinate offsets calculated from two manually selected reference vessel centers. This approach identified 60 vessels across seven ventricular samples for subsequent vessel-centered spatial analyses.

#### Vessel-centered radial and angular analysis

For each identified coronary vessel, spatial transcript coordinates were transformed into a vessel-centered reference frame to characterize gene expression according to radial distance and angular position relative to the vessel center. Additionally, transcripts located within 400 μm of each vessel center were retained for downstream analysis. To account for differences in vessel geometry, radial distances were normalized to the estimated vessel wall boundary. The boundary was approximated using transcripts present towards the edge of the vessel wall (*Myh11, Tagln, Flna, Fstl1, Itgb1, Smtn, Acta2)* with the mean distance of the three nearest transcripts calculated within successive 5° angular intervals. The radial distance of each transcript was then divided by the estimated vessel wall radius at its corresponding angular position, generating a normalized radial coordinate that accounted for vessel eccentricity.

To identify spatially organized transcriptional programs, Moran’s I spatial autocorrelation was calculated independently for each gene within each vessel using the spatial_autocorr function in Squidpy (v1.8.1) with 100 permutations. The 500 most spatially autocorrelated genes per vessel were retained for subsequent analysis. Radial and angular transcript distributions were represented using 50-bin histograms and normalized to the corresponding global transcript distributions. Genes were then grouped according to similarities in their spatial distributions using Bayesian Gaussian mixture models (BayesianGaussianMixture, scikit-learn, v1.8.0) with a Dirichlet process prior. Separate models were fitted to radial and angular distributions, specifying 20 and 10 components, respectively.

To identify transcriptional programs shared across vessels, gene lists from individual vessel-specific clusters were compared using hypergeometric tests implemented in SciPy (v1.17.1) with the hypergeom function. Statistical significance was assessed following Benjamini–Hochberg false discovery rate correction using the multipletests function with the statsmodels package (v0.14.6). Significant gene-list overlaps were represented as edges in a network in which individual vessel-specific clusters constituted nodes. Related clusters were grouped into metaclusters using the Louvain community detection algorithm implemented in NetworkX (v3.6.1) (louvain_communities, resolution = 2). Representative gene signatures were defined by identifying genes present in at least 40% of nodes within each metacluster and represented in at least two nodes. The 20 most frequently occurring genes were retained for downstream biological interpretation. This approach identified 16 transcriptional metaclusters representing vascular cell populations and biological processes, including smooth muscle, fibroblast-associated remodeling, macrophage-associated programs, cell proliferation, and interferon-stimulated gene (ISG) responses.

To assess spatial relationships between transcriptional programs, bivariate Moran’s *I* was used to evaluate spatial associations between program-associated regions within individual vessels, including ISG, fibrosis, macrophage, and proliferation-associated programs. The following equation was used to calculate bivariate Moran’s *I:*

A positive value indicates that the high-density regions of one module are good predictors of the high-density region of the module it is being compared against, with values ranging from −1 to 1.

#### Quantification of spatial transcriptional programs

To quantify the spatial distribution of transcriptional programs, the perivascular region surrounding each vessel was divided into computational hexagonal bins measuring 10 μm in diameter. Hexagons containing at least three transcripts corresponding to genes within a given metacluster were classified as positive for that transcriptional program. The total area occupied by positive hexagons was used to estimate the spatial coverage of each program within individual vessels. Differences in program-associated area between control and TAC vessels were evaluated using two-sided Wilcoxon rank-sum tests implemented with the mannwhitneyu function in SciPy (v1.17.1). Vessel membership within individual metaclusters was determined by network cluster assignment or significant connectivity to an existing metacluster, together with the presence of at least two genes from its representative gene signature. Differences in the frequency of metacluster occurrence between control and TAC vessels were assessed using one-sided Fisher’s exact tests with the fisher_exact function from the SciPy package (v1.17.1).

## Statistical analysis

Statistical analyses were performed to evaluate differences in myocardial remodeling, perivascular fibrosis, and interferon-stimulated gene expression between experimental groups. For RNAscope and Masson’s trichrome quantifications, comparisons between two experimental groups were performed using unpaired, two-tailed Student’s t-tests, whereas comparisons involving three or more groups were performed using one-way analysis of variance (ANOVA). Post hoc multiple-comparison corrections were applied following ANOVA. Data are presented as Mean ± SEM, as indicated in the figure legends. Sample sizes, statistical tests, and exact P values are reported in the corresponding figure legends.

For spatial transcriptomic analyses, differences in the area occupied by individual transcriptional programs between control and TAC vessels were evaluated using two-sided Wilcoxon rank-sum tests (Mann–Whitney U tests), implemented using the mannwhitneyu function in SciPy. Differences in the frequency of transcriptional program occurrence were assessed using one-sided Fisher’s exact tests (fisher_exact, SciPy). Statistical significance of gene-list overlap during network-based metacluster identification was evaluated using hypergeometric tests followed by Benjamini–Hochberg false discovery rate correction.

## Supporting information

Supplement Figures with Captions

Figures with Captions

## Acknowledgements

We thank J. Santini and M. Erb at the UC San Diego School of Medicine Light Microscopy Core for technical assistance with microscopy and imaging experiments; and T. Taniguchi for the gift of the Irf3fl/fl mice. Microscopy and image analysis was performed at the Nikon Imaging Center at UC San Diego.

## Author contributions

Conceptualization: G.C.B., V.K.N., and K.R.K., Methodology: G.C.B., S.J.W, V.K.N, and K.R.K., Investigation: G.C.B., S.J.W., B.B.G., V.K.N., B.Z., J.M.H., R.P.V., and Z.F. Visualization: G.C.B. and S.J.W. Formal analysis: G.C.B. and S.J.W. Software: S.J.W. Supervision: Z.F. and K.R.K. Data curation: G.C.B. and S.J.W. Validation: Z.F. and K.R.K. Project administration: G.C.B., and K.R.K. Writing—original draft: G.C.B. and S.J.W. Writing—review and editing: G.C.B., S.J.W., B.B.G., V.K.N., B.Z., J.M.H., and K.R.K. Resources: V.K.N., J.M.H., R.P.V., E.G., J.H. Funding acquisition: J.M.H., V.K.N., and K.R.K.

## Data, code, and materials availability

All data and code needed to evaluate and reproduce the results in the paper are present through the paper and/or Supplementary Materials. No new materials were generated in this study. Samples used in this study are maintained by the corresponding authors.

---

**Supplementary Figure 1. Histological assessment of ventricular interstitial collagen deposition following transverse aortic constriction (TAC)**

**(A)** Representative higher-magnification Masson’s trichrome-stained myocardial sections from the left ventricle (LV; top) and right ventricle (RV; bottom) of control and wild-type (WT) mice following transverse aortic constriction (TAC). Collagen deposition is visualized in blue, and myocardial tissue is stained red. Images illustrate the distribution of interstitial collagen within the ventricular myocardium, complementing the quantitative assessment of interstitial fibrosis presented in Fig. 1C–D. Scale bars, 50 μm.

**Supplementary Figure 2. Computational identification of coronary vessels across control and TAC hearts.**

**(A)** Spatial transcriptomic maps of all 60 coronary vascular regions identified across control (n = 13 vessels) and transverse aortic constriction (TAC; n = 47 vessels) samples using the Xenium platform. Each circular region represents an individual vessel-centered spatial transcriptomic field, with smooth muscle-associated transcripts (*Myh11, Tagln, Acta2, and Fstl1*) shown in purple. The spatial organization of these transcripts was used to visualize vascular structures and support computational vessel identification. The identified vessels were subsequently used for radial and angular analyses of perivascular transcriptional programs.

**Supplementary Figure 3. Angular estimation of vessel wall geometry for vessel-centered spatial analysis.**

**(A)** Representative scatterplot illustrating the distribution of smooth muscle-associated transcripts in vessel-centered polar coordinates. Each blue point represents a spatial transcript positioned according to its angular coordinate (θ, 0–360°) and radial distance from the vessel center (μm). The orange line represents the estimated vessel wall radius at each angular position, calculated using the mean distance of the three nearest smooth muscle-associated transcripts within successive 5° angular intervals. This angularly resolved estimate of vessel wall geometry was used to normalize radial transcript distances and account for differences in vessel eccentricity during downstream spatial transcriptomic analyses.

**Supplementary Figure 4. Network-based identification of spatial transcriptional metaclusters across coronary arteries.**

**(A)** Network representation of vessel-specific Gaussian mixture model (GMM) clusters identified from Xenium spatial transcriptomic analysis of 60 coronary vessels. Each node represents an individual vessel-specific GMM cluster, with node colors indicating the vessel of origin. Edges represent significant overlap between cluster-associated gene sets. **(B)** Louvain community detection applied to the network in (A) to identify transcriptional metaclusters shared across vessels. Nodes are colored according to their assigned metacluster, with representative genes shown for major transcriptional programs, including interferon-stimulated genes (ISGs), cardiomyocytes, vascular smooth muscle cells, fibroblasts, proliferating cells, endothelial populations, macrophages, and lymphatic endothelial cells. Metaclusters were annotated based on their representative gene signatures.

**Supplementary Figure 5. Vessel-level distribution of spatial transcriptional programs in control and TAC coronary vessels.**

**(A)** Dot plot illustrating the distribution of spatial transcriptional programs across individual coronary vessels identified in control and transverse aortic constriction (TAC) hearts using Xenium spatial transcriptomics. Each column represents an individual coronary vessel, grouped by experimental condition (control, green; TAC, red), with vessel cross-sectional area (μm²) displayed in the upper bar plot. Rows represent transcriptional programs associated with fibrosis, vascular smooth muscle, macrophages, lymphatic endothelial cells, cardiomyocytes, proliferation, vascular endothelial cells, B cells, interferon-stimulated genes (ISGs), and T cells. Dot size indicates the relative area occupied by each transcriptional program above its enrichment threshold, while dot color represents the degree of enrichment relative to the corresponding background or reference distribution. This analysis illustrates vessel-to-vessel heterogeneity in the spatial representation of transcriptional programs and provides an overview of cellular and molecular remodeling within the coronary perivascular microenvironment following pressure overload.

**Supplementary Figure 6. Transverse aortic constriction induces *Nppa* expression in both coronary vascular territories.**

**(A)** Representative whole-heart RNAscope images showing *Nppa* expression (cyan) in control and wild-type (WT) mice following transverse aortic constriction (TAC). **(B)** Quantification of *Nppa*-positive area in the left coronary artery (LCA) and right coronary artery (RCA) territories, expressed as a percentage of total tissue area. TAC significantly increased *Nppa* expression in both coronary territories compared with controls. Data are presented as mean ± SEM, with individual data points representing individual analyzed regions. Statistical analyses were performed using unpaired, two-tailed Student’s t-tests. **P < 0.01, ****P < 0.0001.

**Supplementary Figure 7. Proposed model of cGAS–IRF3–IFNAR signaling in the perivascular interferon response.**

**(A)** Schematic illustrating a proposed mechanism linking mechanical stress to localized interferon-stimulated gene (ISG) expression in the coronary perivascular microenvironment. Pressure overload-associated mechanical stress is hypothesized to promote activation of the cGAS–STING–IRF3 pathway in an initiating cell, potentially leading to type I interferon (IFN-α/β) production and secretion. Secreted type I interferons may subsequently activate IFNAR signaling in neighboring responder cells, promoting downstream JAK–STAT signaling and ISG expression. Genetic deletion of CGAS or IRF3 and pharmacological blockade of IFNAR attenuated perivascular Ifit1 expression following transverse aortic constriction (TAC). The initiating cell type, source of cytosolic DNA, involvement of STING, and cellular origin of type I interferons remain to be established.

**Supplementary Figure 8. Genetic deletion of IRF3 or cGAS does not significantly alter *Nppa* expression following TAC.**

**(A)** Quantification of *Nppa*-positive area surrounding the left coronary artery (LCA) and right coronary artery (RCA) in wild-type (WT), Irf3^−/−^, and Cgas^−/−^ mice following transverse aortic constriction (TAC), expressed as a percentage of the analyzed perivascular area. No statistically significant differences in perivascular *Nppa* expression were detected among genotypes in either coronary territory. **(B)** Quantification of *Nppa*-positive area within the left ventricular (LV) and right ventricular (RV) myocardium, expressed as a percentage of the total analyzed tissue area. No statistically significant differences in myocardial *Nppa* expression were detected among WT, Irf3^−/−^, and Cgas^−/−^ TAC mice in either ventricular region. Data are presented as mean ± SEM, with individual data points representing individual analyzed regions Statistical analyses were performed using one-way ANOVA. ns, not significant.

**Supplementary Figure 9. Genetic deletion of IRF3 or cGAS does not significantly alter early perivascular fibrosis following TAC.**

**(A)** Representative whole-heart Masson’s trichrome-stained sections from wild-type (WT), Irf3^−/−^, and Cgas^−/−^ mice following transverse aortic constriction (TAC), illustrating myocardial collagen deposition. **(B)** Representative higher-magnification Masson’s trichrome images showing collagen deposition surrounding the left coronary artery (LCA; top) and right coronary artery (RCA; bottom) in WT, Irf3^−/−^, and Cgas^−/−^ TAC hearts. Collagen is stained blue, and myocardial tissue is stained red. Scale bars, 50 μm. **(C–D)** Quantification of perivascular collagen-positive area surrounding the LCA (C) and RCA (D), expressed as a percentage of the perivascular area. No statistically significant differences in perivascular collagen deposition were detected among WT, Irf3^−/−^, and Cgas^−/−^ TAC groups in either coronary territory. Data are presented as mean ± SEM, with individual data points representing individual vessels. Statistical analyses were performed using one-way ANOVA. ns, not significant.

**Supplementary Figure 10. IFNAR blockade does not significantly alter perivascular or myocardial *Nppa* expression following TAC.**

**(A)** Representative whole-heart RNAscope images showing *Nppa* (cyan) and *Ifit1* (magenta) expression in wild-type mice treated with isotype control or anti-IFNAR antibody following transverse aortic constriction (TAC). **(B)** Quantification of *Nppa*-positive area within the left ventricular (LV) and right ventricular (RV) myocardium, expressed as a percentage of total analyzed tissue area. No statistically significant differences in myocardial *Nppa* expression were detected between treatment groups in either ventricular region. Data are presented as mean ± SEM, with individual data points representing individual analyzed regions. Statistical analyses were performed using unpaired, two-tailed Student’s t-tests. ns, not significant.

**Supplementary Figure 11. IFNAR blockade does not significantly reduce ventricular interstitial fibrosis following TAC.**

**(A)** Representative whole-heart Masson’s trichrome-stained sections from wild-type mice treated with isotype control or anti-IFNAR antibody following transverse aortic constriction (TAC). Collagen deposition is visualized in blue, and myocardial tissue is stained red. **(B)** Quantification of interstitial collagen-positive area in the left ventricular (LV) and right ventricular (RV) myocardium, expressed as a percentage of total myocardial area. No statistically significant differences in interstitial collagen deposition were detected between isotype control- and anti-IFNAR-treated TAC mice in either ventricular region. Data are presented as mean ± SEM, with individual data points representing individual analyzed regions. Statistical analyses were performed using unpaired, two-tailed Student’s t-tests. ns, not significant.

**Supplementary Figure 12. Perivascular collagen deposition following IFNAR blockade in TAC mice.**

**(A–B)** Representative Masson’s trichrome-stained sections and quantification of perivascular collagen-positive area in the left coronary artery (LCA; A) and right coronary artery (RCA; B) territories of mice subjected to transverse aortic constriction (TAC) and treated with isotype control or anti-IFNAR antibody. Collagen-positive area was quantified as a percentage of the total perivascular area. Each dot represents individual vessel. Data are presented as mean ± SEM. ns, not significant. Scale bars, 50 μm.

