## Supplement Figures with Captions for "Acute hypertension causes focal coronary artery perivascular cGAS-, IRF3, and IFNAR-dependent type I interferon responses"

*Gilda Castellanos von Borstel*[1], *Samuel J. Wright*[1,2], *Betul B. Gunes*[3], *Van K. Ninh*[3], *Bo Zhang*[1], *Jessica M. Hong*[3], *Raimon Padrós-Valls*[1,2], *Eugene Golts*[4], Jingjing Hu[5], Zhenxing Fu[2], *Kevin R. King[1,2,3,]

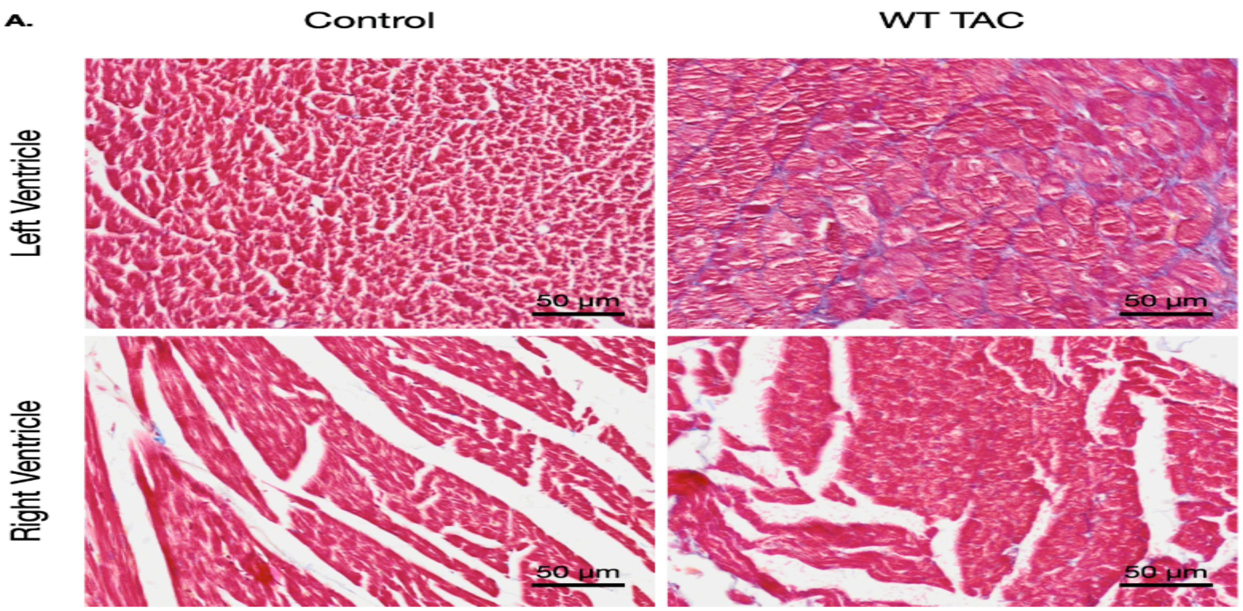


Supp fig 1.

**Supplementary Figure 1. Histological assessment of ventricular interstitial collagen deposition following transverse aortic constriction (TAC)**

Representative higher-magnification Masson's trichrome-stained myocardial sections from the left ventricle (LV; top) and right ventricle (RV; bottom) of control and wild-type (WT) mice following transverse aortic constriction (TAC). Collagen deposition is visualized in blue, and myocardial tissue is stained red. Images illustrate the distribution of interstitial collagen within the ventricular myocardium, complementing the quantitative assessment of interstitial fibrosis presented in Fig. 1C–D. Scale bars, 50 μm.

**A. Smooth muscle-associated transcripts across 60 vessels**

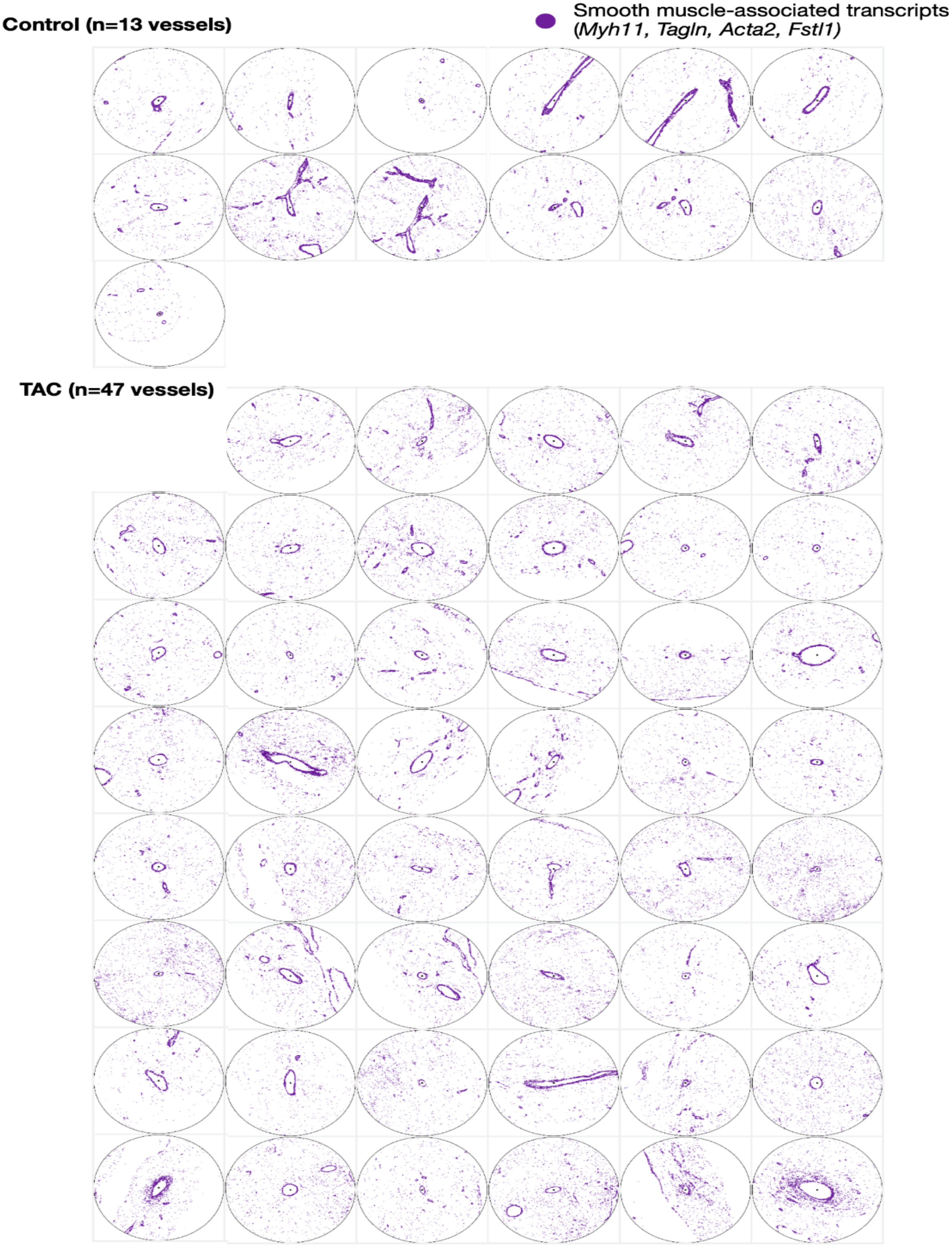

**Supplementary Figure 2. Computational identification of coronary vessels across control and TAC hearts.**

Spatial transcriptomic maps of all 60 coronary vascular regions identified across control (n = 13 vessels) and transverse aortic constriction (TAC; n = 47 vessels) samples using the Xenium platform. Each circular region represents an individual vessel-centered spatial transcriptomic field, with smooth muscle-associated transcripts (*Myh11, Tagln, Acta2, and Fstl1*) shown in purple. The spatial organization of these transcripts was used to visualize vascular structures and support computational vessel identification. The identified vessels were subsequently used for radial and angular analyses of perivascular transcriptional programs.

A.

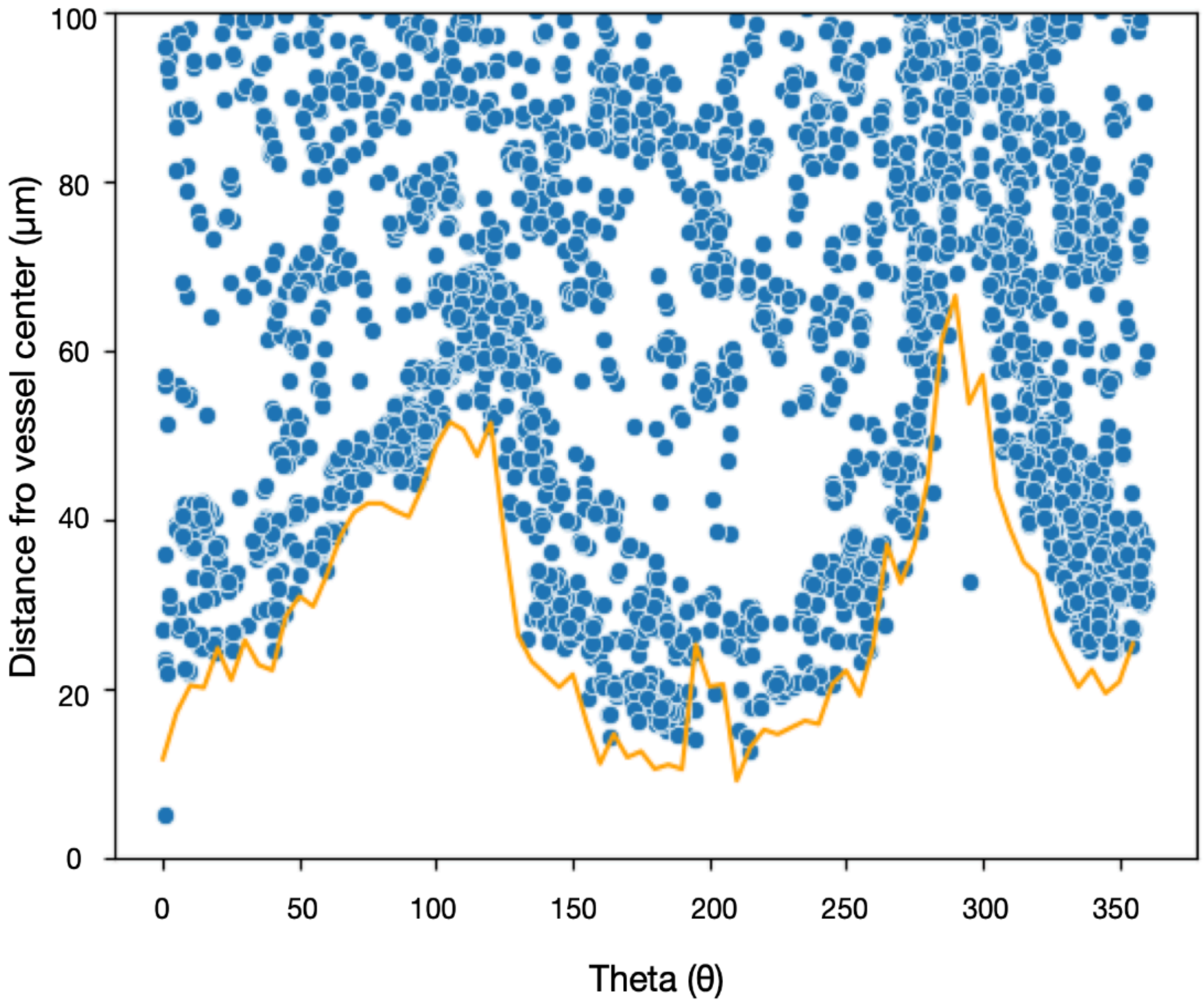


**Supplementary Figure 3. Angular estimation of vessel wall geometry for vessel-centered spatial analysis.**

Representative scatterplot illustrating the distribution of smooth muscle-associated transcripts in vessel-centered polar coordinates. Each blue point represents a spatial transcript positioned according to its angular coordinate (θ, 0–360°) and radial distance from the vessel center (µm). The orange line represents the estimated vessel wall radius at each angular position, calculated using the mean distance of the three nearest smooth muscle-associated transcripts within successive 5° angular intervals. This angularly resolved estimate of vessel wall geometry was used to normalize radial transcript distances and account for differences in vessel eccentricity during downstream spatial transcriptomic analyses.

**Supp fig 4.**

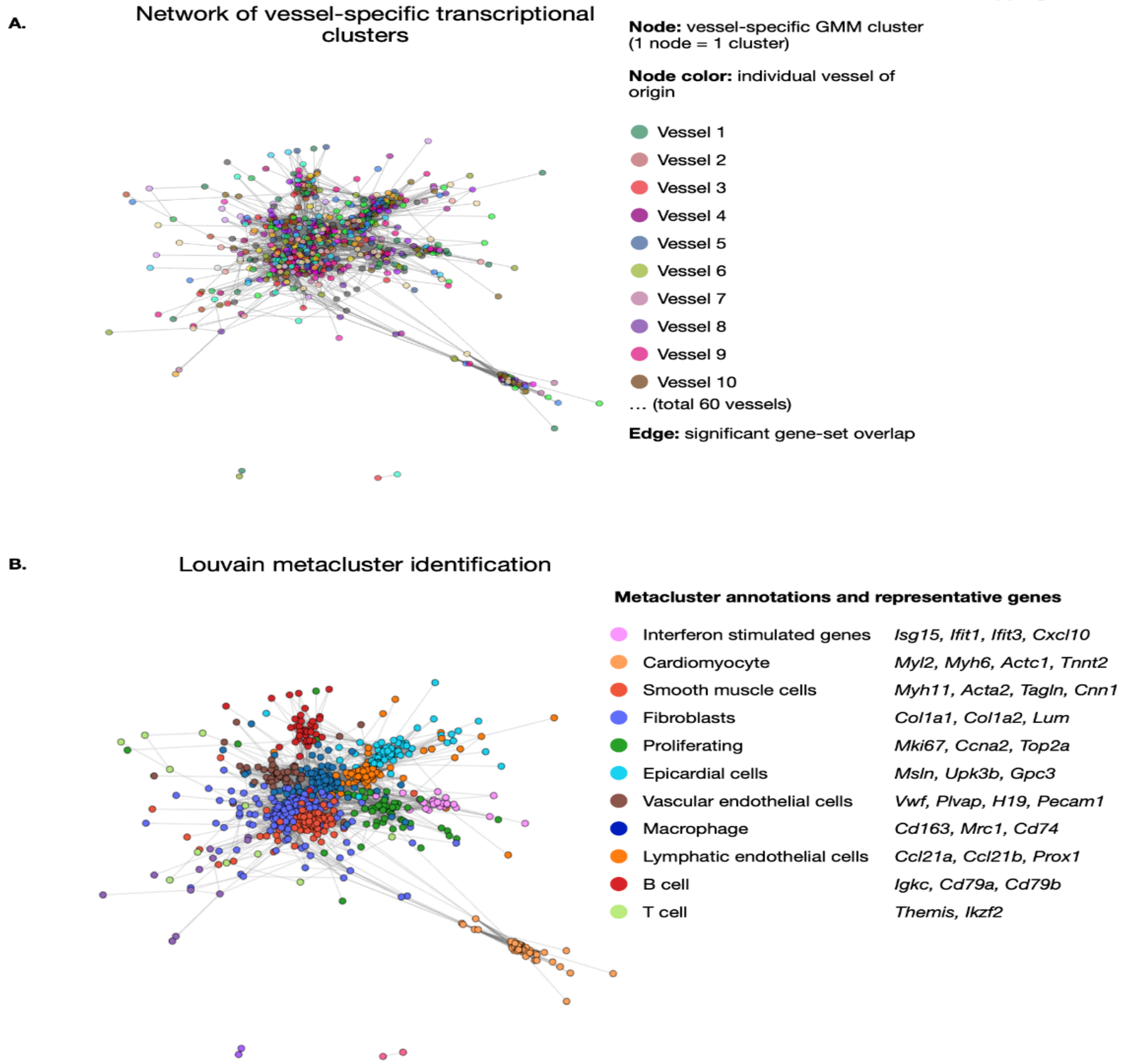


**Supplementary Figure 4. Network-based identification of spatial transcriptional metaclusters across coronary arteries.**

(**A**) Network representation of vessel-specific Gaussian mixture model (GMM) clusters identified from Xenium spatial transcriptomic analysis of 60 coronary vessels. Each node represents an individual vessel-specific GMM cluster, with node colors indicating the vessel of origin. Edges represent significant overlap between cluster-associated gene sets. (**B**) Louvain community detection applied to the network in (A) to identify transcriptional metaclusters shared across vessels. Nodes are colored according to their assigned metacluster, with representative genes shown for major transcriptional programs, including interferon-stimulated genes (ISGs), cardiomyocytes, vascular smooth muscle cells, fibroblasts, proliferating cells, endothelial populations, macrophages, and lymphatic endothelial cells. Metaclusters were annotated based on their representative gene signatures.

Supp fig 5.

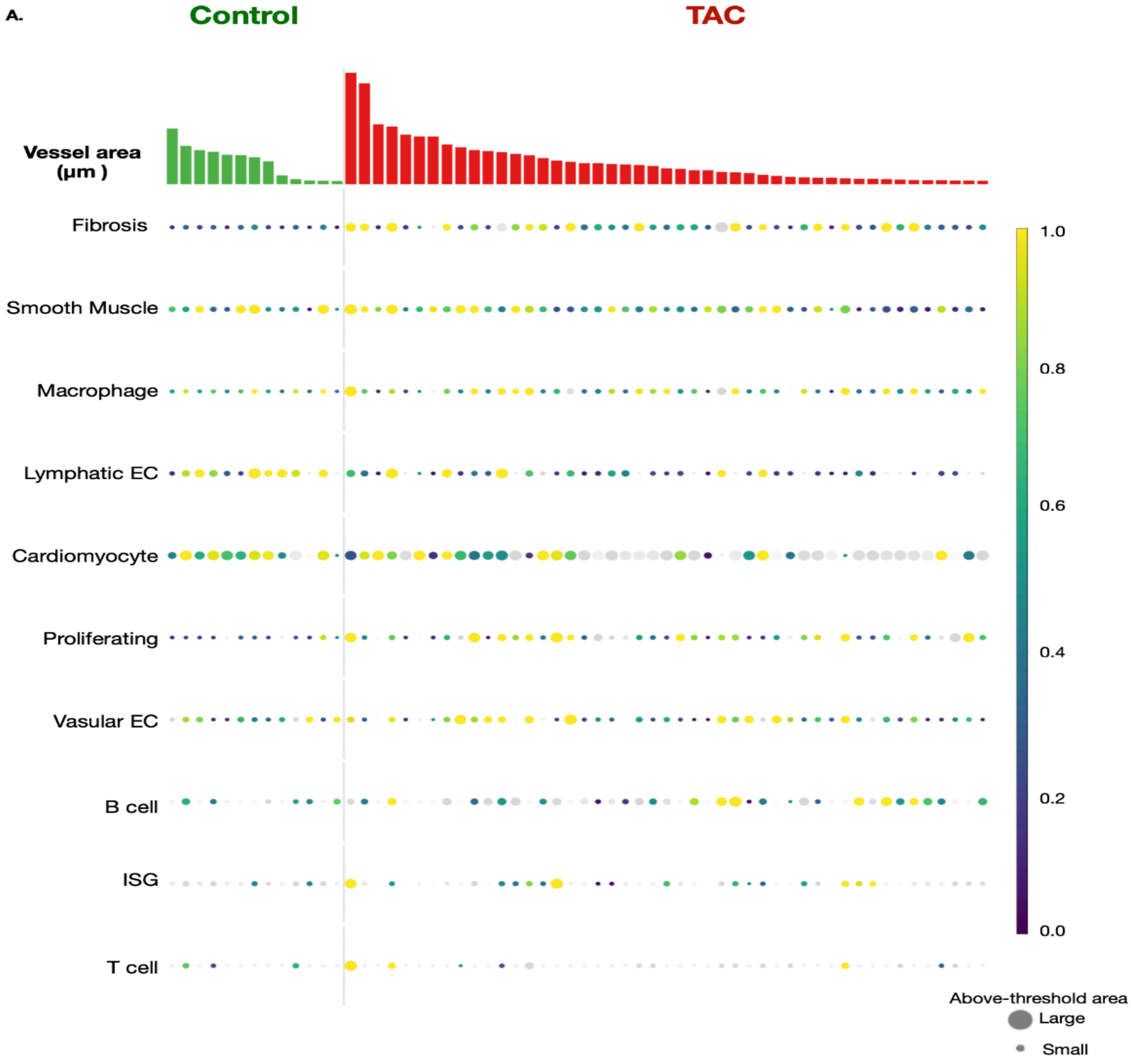


**Supplementary Figure 5. Vessel-level distribution of spatial transcriptional programs in control and TAC coronary vessels.**

Dot plot illustrating the distribution of spatial transcriptional programs across individual coronary vessels identified in control and transverse aortic constriction (TAC) hearts using Xenium spatial transcriptomics. Each column represents an individual coronary vessel, grouped by experimental condition (control, green; TAC, red), with vessel cross-sectional area (µm²) displayed in the upper bar plot. Rows represent transcriptional programs associated with fibrosis, vascular smooth muscle, macrophages, lymphatic endothelial cells, cardiomyocytes, proliferation, vascular endothelial cells, B cells, interferon-stimulated genes (ISGs), and T cells. Dot size indicates the relative area occupied by each transcriptional program above its enrichment threshold, while dot color represents the degree of enrichment relative to the corresponding background or reference distribution. This analysis illustrates vessel-to-vessel heterogeneity in the spatial representation of transcriptional programs and provides an overview of cellular and molecular remodeling within the coronary perivascular microenvironment following pressure overload.

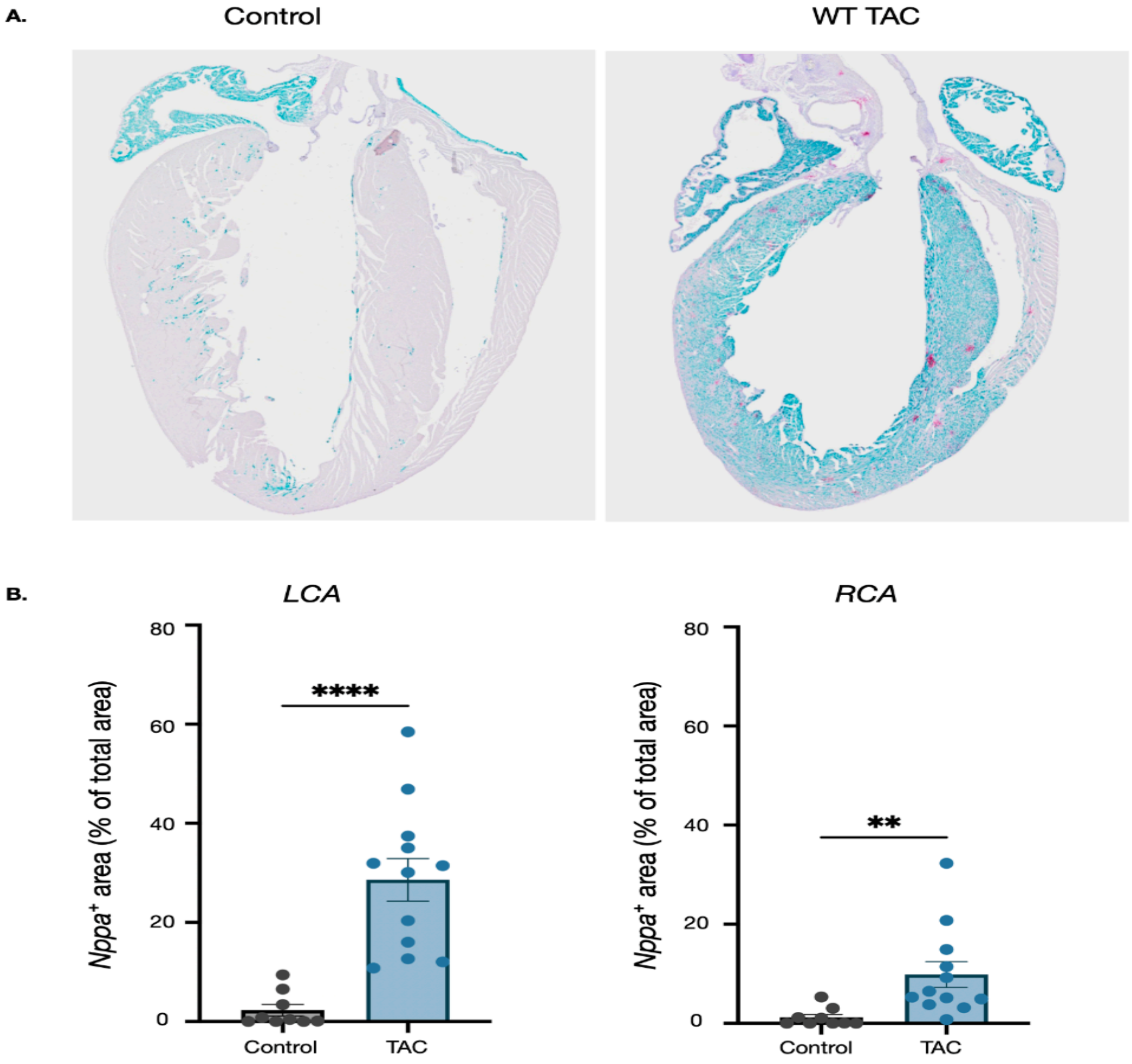


**Supplementary Figure 6. Transverse aortic constriction induces *Nppa* expression in both coronary vascular territories.**

(**A**) Representative whole-heart RNAscope images showing *Nppa* expression (cyan) in control and wild-type (WT) mice following transverse aortic constriction (TAC). (**B**) Quantification of *Nppa*-positive area in the left coronary artery (LCA) and right coronary artery (RCA) territories, expressed as a percentage of total tissue area. TAC significantly increased *Nppa* expression in both coronary territories compared with controls. Data are presented as mean ± SEM, with individual data points representing individual analyzed regions. Statistical analyses were performed using unpaired, two-tailed Student's t-tests. **P < 0.01, ****P < 0.0001.

A.

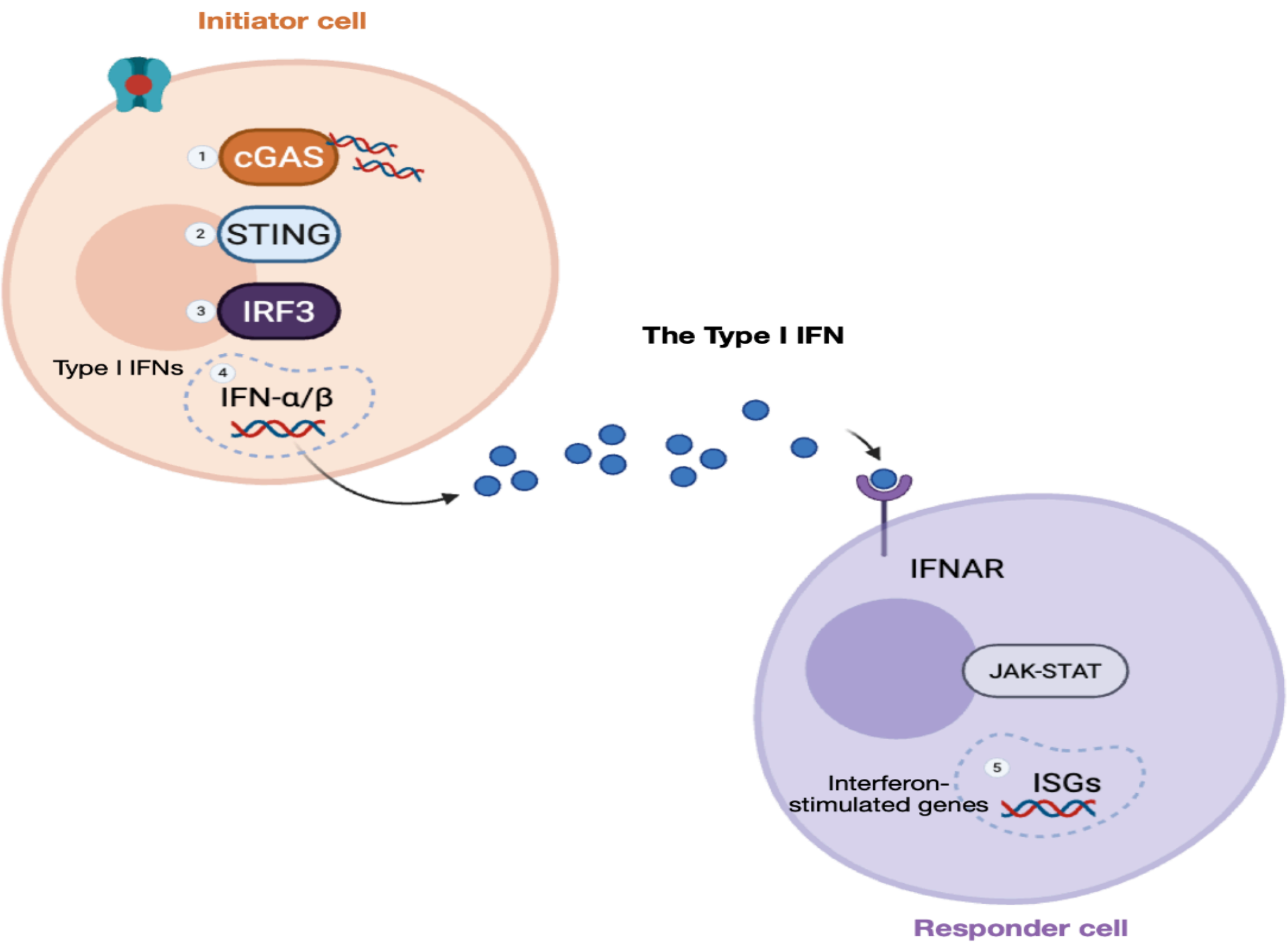


**Supplementary Figure 7. Proposed model of cGAS–IRF3–IFNAR signaling in the perivascular interferon response.**

Schematic illustrating a proposed mechanism linking mechanical stress to localized interferon-stimulated gene (ISG) expression in the coronary perivascular microenvironment. Pressure overload-associated mechanical stress is hypothesized to promote activation of the cGAS–STING–IRF3 pathway in an initiating cell, potentially leading to type I interferon (IFN-α/β) production and secretion. Secreted type I interferons may subsequently activate IFNAR signaling in neighboring responder cells, promoting downstream JAK–STAT signaling and ISG expression. Genetic deletion of CGAS or IRF3 and pharmacological blockade of IFNAR attenuated perivascular Ifit1 expression following transverse aortic constriction (TAC). The initiating cell type, source of cytosolic DNA, involvement of STING, and cellular origin of type I interferons remain to be established.

Supp fig 8.

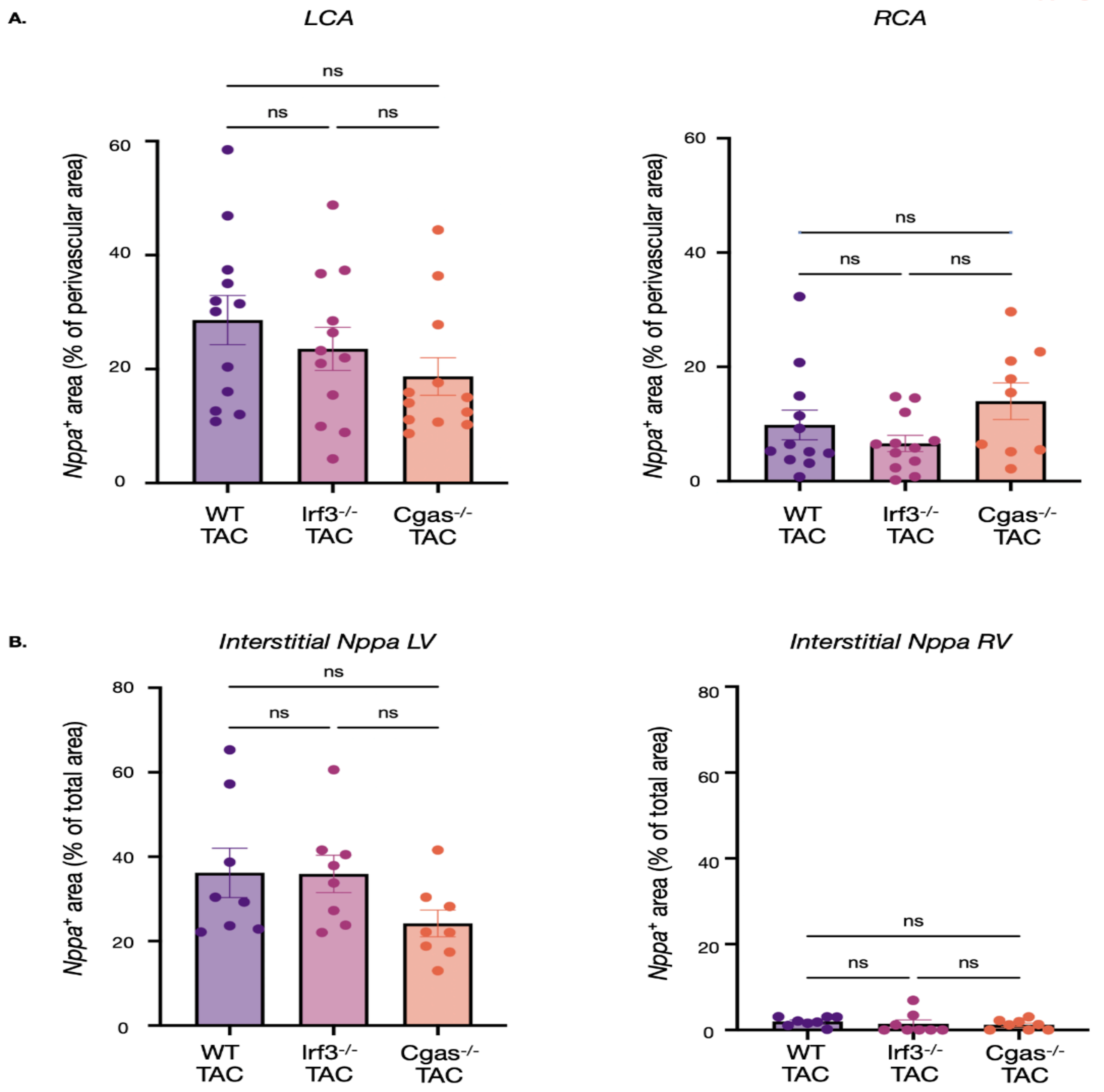

A.
LCA
RCA
Nppa+ area (% of perivascular area)
ns
0
20
40
60
WT TAC
Irf3-/- TAC
Cgas-/- TAC
B.
Interstitial Nppa LV
Interstitial Nppa RV
Nppa+ area (% of total area)
80

**Supplementary Figure 8. Genetic deletion of IRF3 or cGAS does not significantly alter *Nppa* expression following TAC.**

(**A**) Quantification of *Nppa*-positive area surrounding the left coronary artery (LCA) and right coronary artery (RCA) in wild-type (WT), Irf3$^{-/-}$, and Cgas$^{-/-}$ mice following transverse aortic constriction (TAC), expressed as a percentage of the analyzed perivascular area. No statistically significant differences in perivascular *Nppa* expression were detected among genotypes in either coronary territory. (**B**) Quantification of *Nppa*-positive area within the left ventricular (LV) and right ventricular (RV) myocardium, expressed as a percentage of the total analyzed tissue area. No statistically significant differences in myocardial *Nppa* expression were detected among WT, Irf3$^{-/-}$, and Cgas$^{-/-}$ TAC mice in either ventricular region. Data are presented as mean ± SEM, with individual data points representing individual analyzed regions Statistical analyses were performed using one-way ANOVA. ns, not significant.

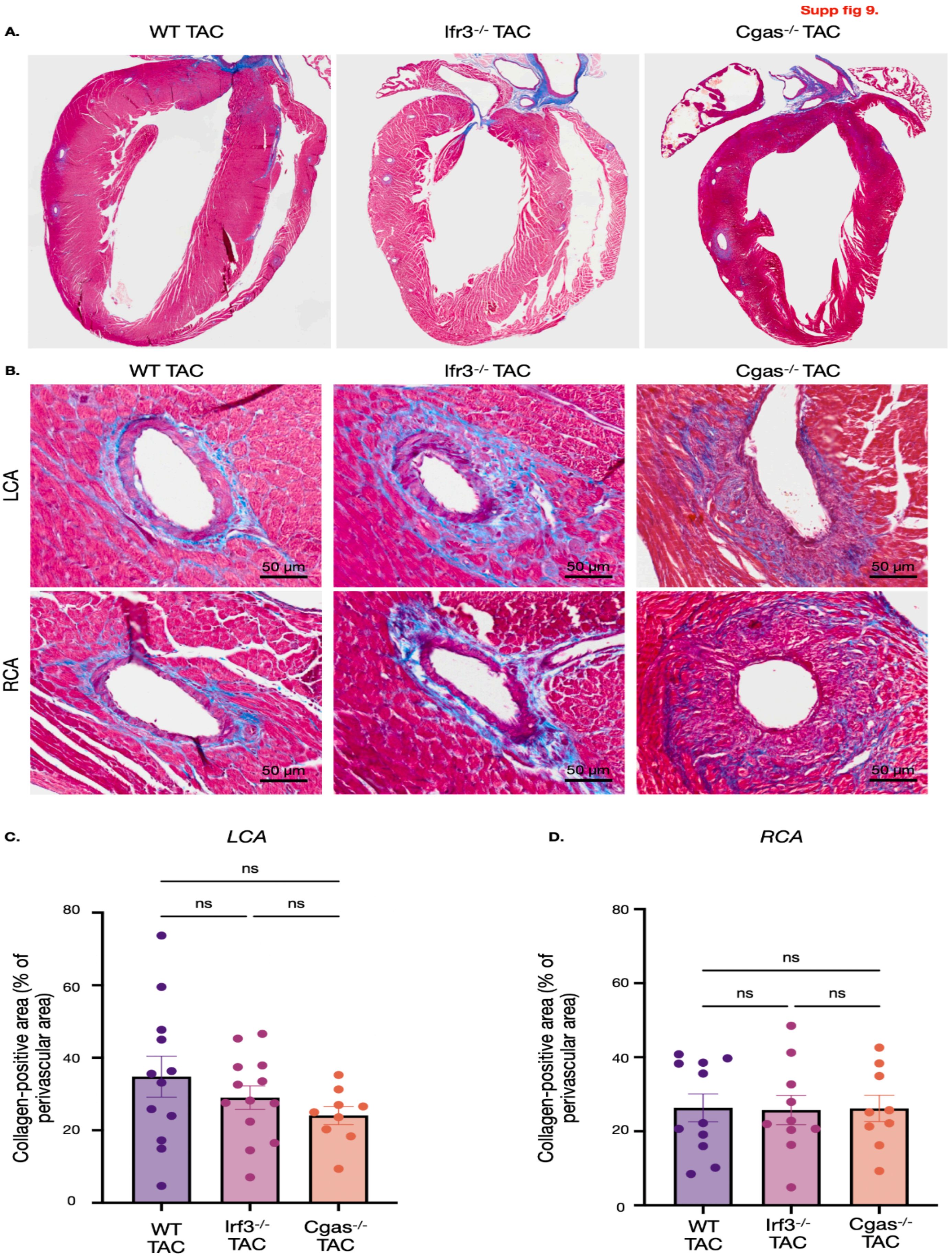

Supp fig 9.
A.
WT TAC
Ifr3-/- TAC
Cgas-/- TAC
B.
WT TAC
Ifr3-/- TAC
Cgas-/- TAC
LCA
RCA
50 μm
C.
LCA
ns
Collagen-positive area (% of perivascular area)
0
20
40
60
80
WT TAC
Irf3-/- TAC
Cgas-/- TAC
D.
RCA
ns
Collagen-positive area (% of perivascular area)
0
20
40
60
80
WT TAC
Irf3-/- TAC
Cgas-/- TAC

**Supplementary Figure 9. Genetic deletion of IRF3 or cGAS does not significantly alter early perivascular fibrosis following TAC.**

(**A**) Representative whole-heart Masson's trichrome-stained sections from wild-type (WT), $Irf3^{-/-}$, and $Cgas^{-/-}$ mice following transverse aortic constriction (TAC), illustrating myocardial collagen deposition. (**B**) Representative higher-magnification Masson's trichrome images showing collagen deposition surrounding the left coronary artery (LCA; top) and right coronary artery (RCA; bottom) in WT, $Irf3^{-/-}$, and $Cgas^{-/-}$ TAC hearts. Collagen is stained blue, and myocardial tissue is stained red. Scale bars, 50 μm. (**C–D**) Quantification of perivascular collagen-positive area surrounding the LCA (C) and RCA (D), expressed as a percentage of the perivascular area. No statistically significant differences in perivascular collagen deposition were detected among WT, $Irf3^{-/-}$, and $Cgas^{-/-}$ TAC groups in either coronary territory. Data are presented as mean ± SEM, with individual data points representing individual vessels. Statistical analyses were performed using one-way ANOVA. ns, not significant.

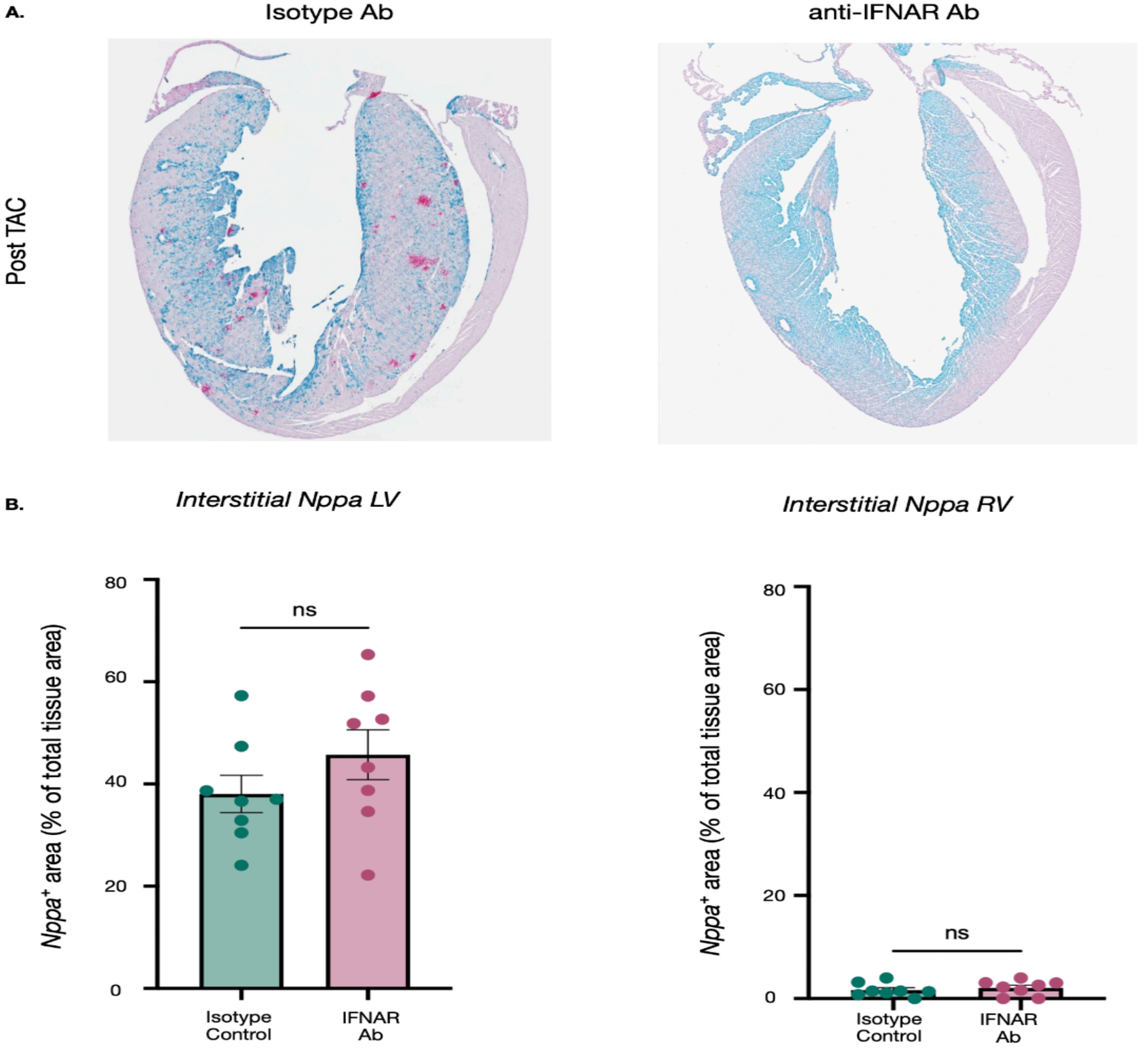
A.
Isotype Ab
anti-IFNAR Ab
Post TAC
B.
Interstitial Nppa LV
Interstitial Nppa RV
Nppa+ area (% of total tissue area)
80
60
40
20
0
ns
Isotype Control
IFNAR Ab

**Supplementary Figure 10. IFNAR blockade does not significantly alter perivascular or myocardial *Nppa* expression following TAC.**

(**A**) Representative whole-heart RNAscope images showing *Nppa* (cyan) and *Ifit1* (magenta) expression in wild-type mice treated with isotype control or anti-IFNAR antibody following transverse aortic constriction (TAC). (**B**) Quantification of *Nppa*-positive area within the left ventricular (LV) and right ventricular (RV) myocardium, expressed as a percentage of total analyzed tissue area. No statistically significant differences in myocardial *Nppa* expression were detected between treatment groups in either ventricular region. Data are presented as mean ± SEM, with individual data points representing individual analyzed regions. Statistical analyses were performed using unpaired, two-tailed Student's t-tests. ns, not significant.

Supp fig 11.

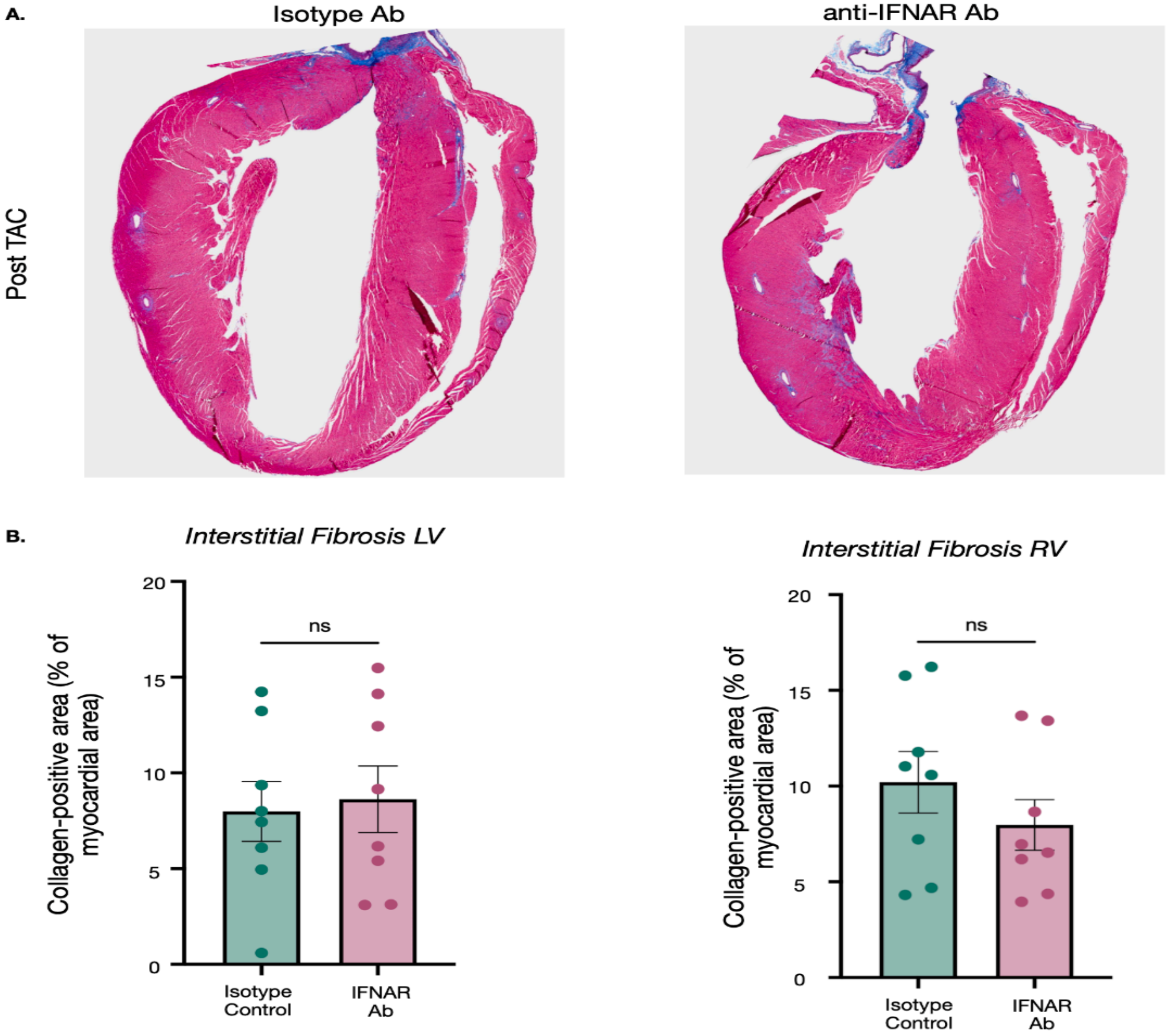
A.
Isotype Ab
anti-IFNAR Ab
Post TAC
B.
Interstitial Fibrosis LV
Interstitial Fibrosis RV
Collagen-positive area (% of myocardial area)
ns
20
15
10
5
0
Isotype Control
IFNAR Ab

**Supplementary Figure 11. IFNAR blockade does not significantly reduce ventricular interstitial fibrosis following TAC.**

(**A**) Representative whole-heart Masson's trichrome-stained sections from wild-type mice treated with isotype control or anti-IFNAR antibody following transverse aortic constriction (TAC). Collagen deposition is visualized in blue, and myocardial tissue is stained red. (**B**) Quantification of interstitial collagen-positive area in the left ventricular (LV) and right ventricular (RV) myocardium, expressed as a percentage of total myocardial area. No statistically significant differences in interstitial collagen deposition were detected between isotype control- and anti-IFNAR-treated TAC mice in either ventricular region. Data are presented as mean ± SEM, with individual data points representing individual analyzed regions. Statistical analyses were performed using unpaired, two-tailed Student's t-tests. ns, not significant.

**Supp fig 12.**

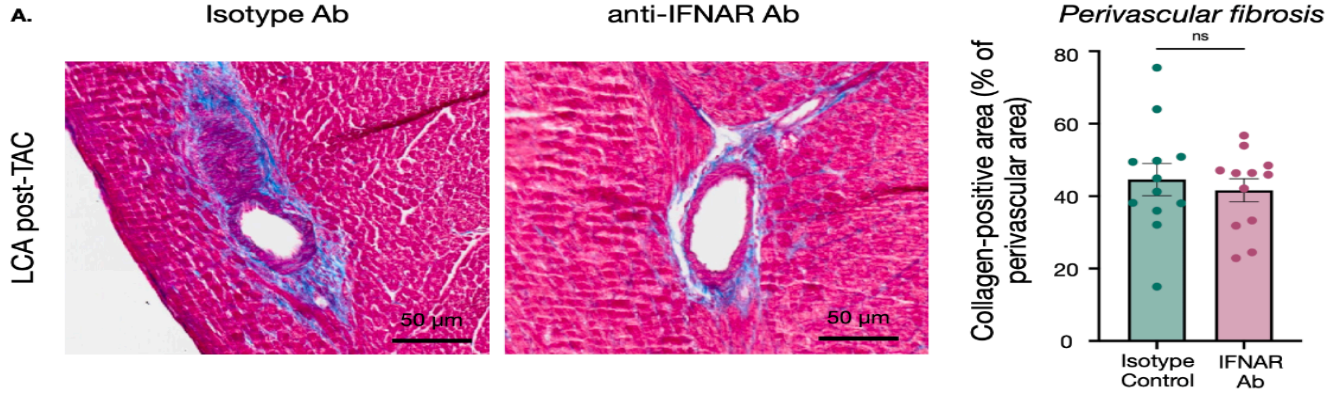

A.
Isotype Ab
anti-IFNAR Ab
Perivascular fibrosis
LCA post-TAC
50 µm
50 µm
Collagen-positive area (% of perivascular area)
80
60
40
20
0
ns
Isotype Control
IFNAR Ab


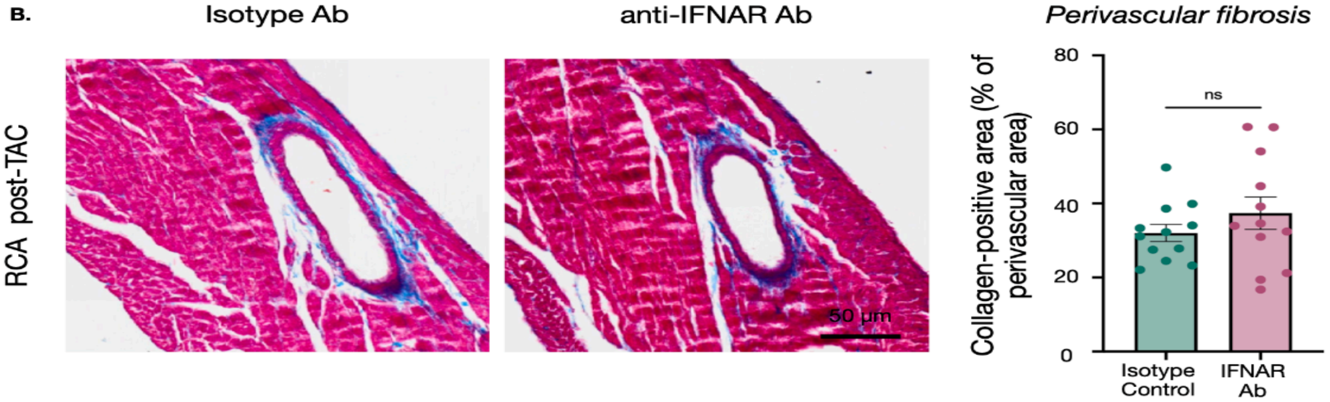

B.
Isotype Ab
anti-IFNAR Ab
Perivascular fibrosis
RCA post-TAC
50 µm
Collagen-positive area (% of perivascular area)
80
60
40
20
0
ns
Isotype Control
IFNAR Ab

**Supplementary Figure 12. Perivascular collagen deposition following IFNAR blockade in TAC mice.** (**A–B**) Representative Masson's trichrome-stained sections and quantification of perivascular collagen-positive area in the left coronary artery (LCA; A) and right coronary artery (RCA; B) territories of mice subjected to transverse aortic constriction (TAC) and treated with isotype control or anti-IFNAR antibody. Collagen-positive area was quantified as a percentage of the total perivascular area. Each dot represents individual vessel. Data are presented as mean ± SEM. ns, not significant. Scale bars, 50 μm.
