## Supplementary material for "Acute hypertension causes focal coronary artery perivascular cGAS-, IRF3, and IFNAR-dependent type I interferon responses": Figures with Captions

Figure 1.

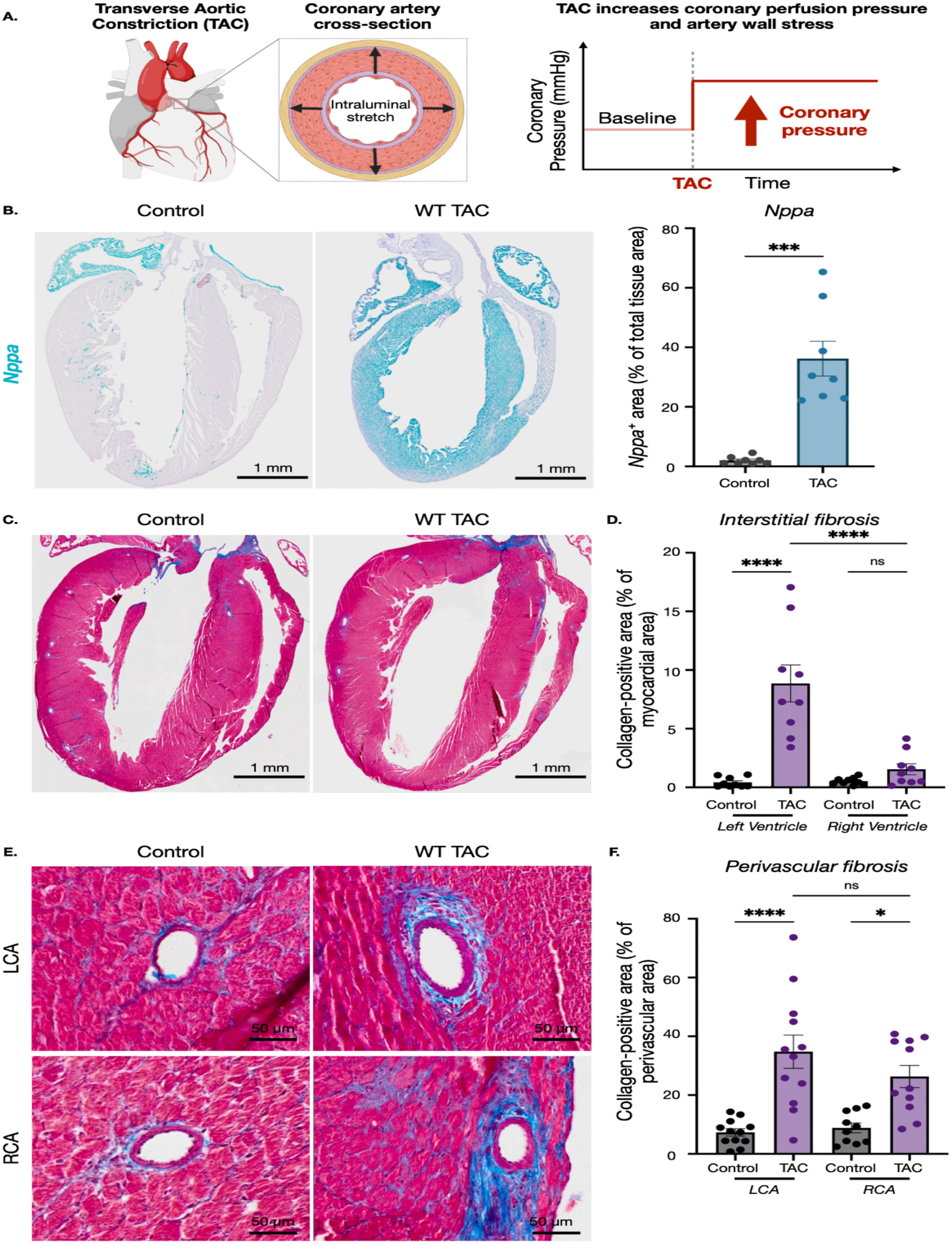

**Figure 1. Hypertensive stress induces bilateral coronary artery perivascular fibrosis**

(**A**) Schematic illustration of transverse aortic constriction (TAC) model used to induce mechanical stress on the myocardium and coronary vasculature through pressure overload. TAC increases proximal aortic and coronary arterial pressure, resulting in increased luminal stress on the coronary vessel wall. (**B**) Representative whole-heart RNAscope images showing *Nppa* expression in control and wild-type (WT) TAC hearts, with corresponding quantifications of *Nppa*-positive area expressed as percentage of total tissue area. Scale bar, 1 mm. **(C)** Whole-heart Masson's trichrome-stained sections from control and WT TAC hearts illustrating increased myocardial collagen deposition (blue) after imposed stress. Scale bar, 1 mm. (**D**) Quantification of interstitial collagen-positive area in the left ventricle (LV) and right ventricle (RV), expressed as a percentage of myocardial area. (**E**) Representative high-magnification Masson's trichrome-stained images of left coronary artery (LCA) and right coronary artery (RCA) from control and WT TAC hearts. Scale bars, 50 µm. (**F**) Quantification of perivascular collagen-positive area surrounding the LCA and RCA, expressed as a percentage of the perivascular area. Bars represent mean ± SEM, with individual data points representing individual arteries. Statistical comparisons were performed using unpaired, two-tailed Student's t-test or one-way ANOVA, as appropriate. Statistical significance is indicated as **P < 0.001, ***P < 0.0001, and ns as non-significant. Exact sample sizes and statistical details are provided in the corresponding Methods section.

Figure 2.

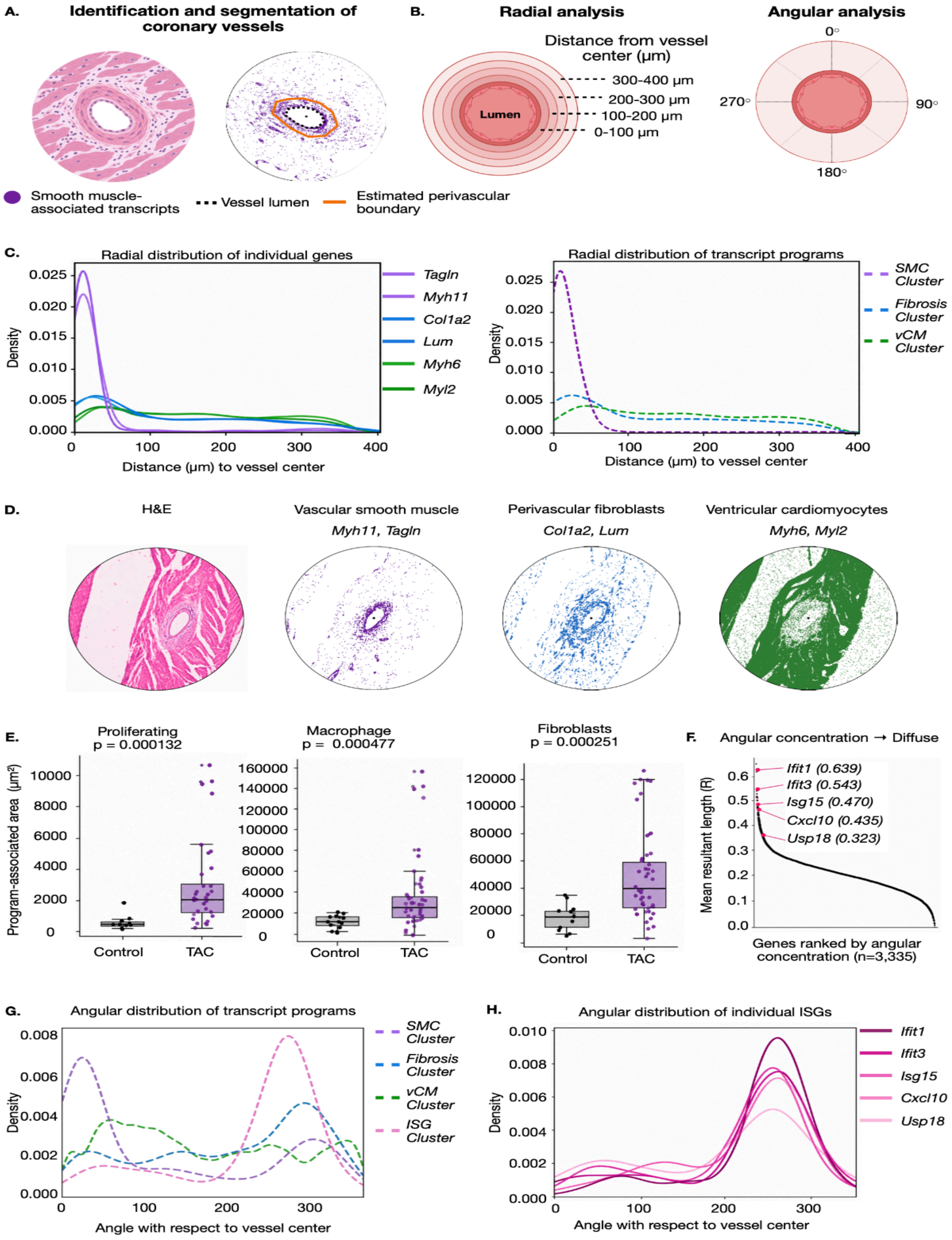
A.
Identification and segmentation of coronary vessels
Smooth muscle-associated transcripts
Vessel lumen
Estimated perivascular boundary
B.
Radial analysis
Distance from vessel center (μm)
Lumen
300-400 μm
200-300 μm
100-200 μm
0-100 μm
Angular analysis
0°
90°
180°
270°
C.
Radial distribution of individual genes
Density
Distance (μm) to vessel center
Tagln
Myh11
Col1a2
Lum
Myh6
Myl2
Radial distribution of transcript programs
SMC Cluster
Fibrosis Cluster
vCM Cluster
D.
H&E
Vascular smooth muscle
Myh11, Tagln
Perivascular fibroblasts
Col1a2, Lum
Ventricular cardiomyocytes
Myh6, Myl2
E.
Proliferating
p = 0.000132
Macrophage
p = 0.000477
Fibroblasts
p = 0.000251
Program-associated area (μm²)
Control
TAC
F.
Angular concentration → Diffuse
Mean resultant length (R)
Ifit1 (0.639)
Ifit3 (0.543)
Isg15 (0.470)
Cxcl10 (0.435)
Usp18 (0.323)
Genes ranked by angular concentration (n=3,335)
G.
Angular distribution of transcript programs
SMC Cluster
Fibrosis Cluster
vCM Cluster
ISG Cluster
Angle with respect to vessel center
H.
Angular distribution of individual ISGs
Ifit1
Ifit3
Isg15
Cxcl10
Usp18
Angle with respect to vessel center

**Figure 2. Vessel-centered spatial transcriptomics reveals focal ISG niches surrounding coronary arteries**

(**A**) Schematic representation of the computational approach used to identify and segment coronary vessels in spatial transcriptomic datasets. Coronary vessels were identified using vascular smooth muscle-associated transcripts, followed by segmentation of the vessel lumen and estimation of the surrounding perivascular boundary. (**B**) Schematic illustrating vessel-centered radial and angular analyses. Radial analysis quantifies gene expression as a function of distance from the vessel center using concentric 100-μm intervals, while angular analysis evaluates the circumferential distribution of gene expression relative to the vessel center. (**C**) Representative radial distributions of individual genes (left) and transcriptional programs identified by Gaussian mixture modeling (GMM; right). Smooth muscle-associated genes (*Myh11, Tagln*) and the corresponding smooth muscle transcriptional program are concentrated near the vessel center, whereas fibroblast-associated genes (*Col1a2, Lum*) and cardiomyocyte-associated genes (*Myh6, Myl2*) exhibit broader radial distributions. Radial transcriptional programs include smooth muscle (SMC), fibrosis-associated, and ventricular cardiomyocyte-associated programs (vCM). (**D**) Representative spatial maps showing the anatomical localization of vascular smooth muscle (*Myh11, Tagln*), perivascular fibroblast (*Col1a2, Lum*), and ventricular cardiomyocyte (*Myh6, Myl2*) transcriptional signatures relative to the coronary vessel (H&E). (**E**) Quantification of program-associated area occupied by proliferation-, macrophage-, and fibroblast- associated transcriptional programs in control and TAC arteries. P values were calculated using two-sided Mann–Whitney U tests. (**F**) Genome-wide ranking of 3,335 genes according to angular concentration, quantified using mean resultant length (R). Higher R values indicate greater angular concentration, whereas lower values indicate more diffuse circumferential distributions. Interferon-stimulated genes, including *Ifit1, Ifit3, Isg15, Cxcl10, and Usp18*, are among genes exhibiting elevated angular concentration. (**G**) Representative angular distributions of transcriptional programs, including smooth muscle, fibrosis-associated, ventricular cardiomyocyte-associated, and ISG-associated programs. (**H**) Angular distributions of individual interferon-stimulated genes *(Ifit1, Ifit3, Isg15, Cxcl10, and Usp18)* within a representative coronary vessel. Spatial transcriptomic analysis included 60 intramyocardial vessels identified across seven mice (control, n = 2; TAC day 7, n = 2; TAC day 28, n = 3), comprising 13 control and 47 TAC vessels.

Figure 3.

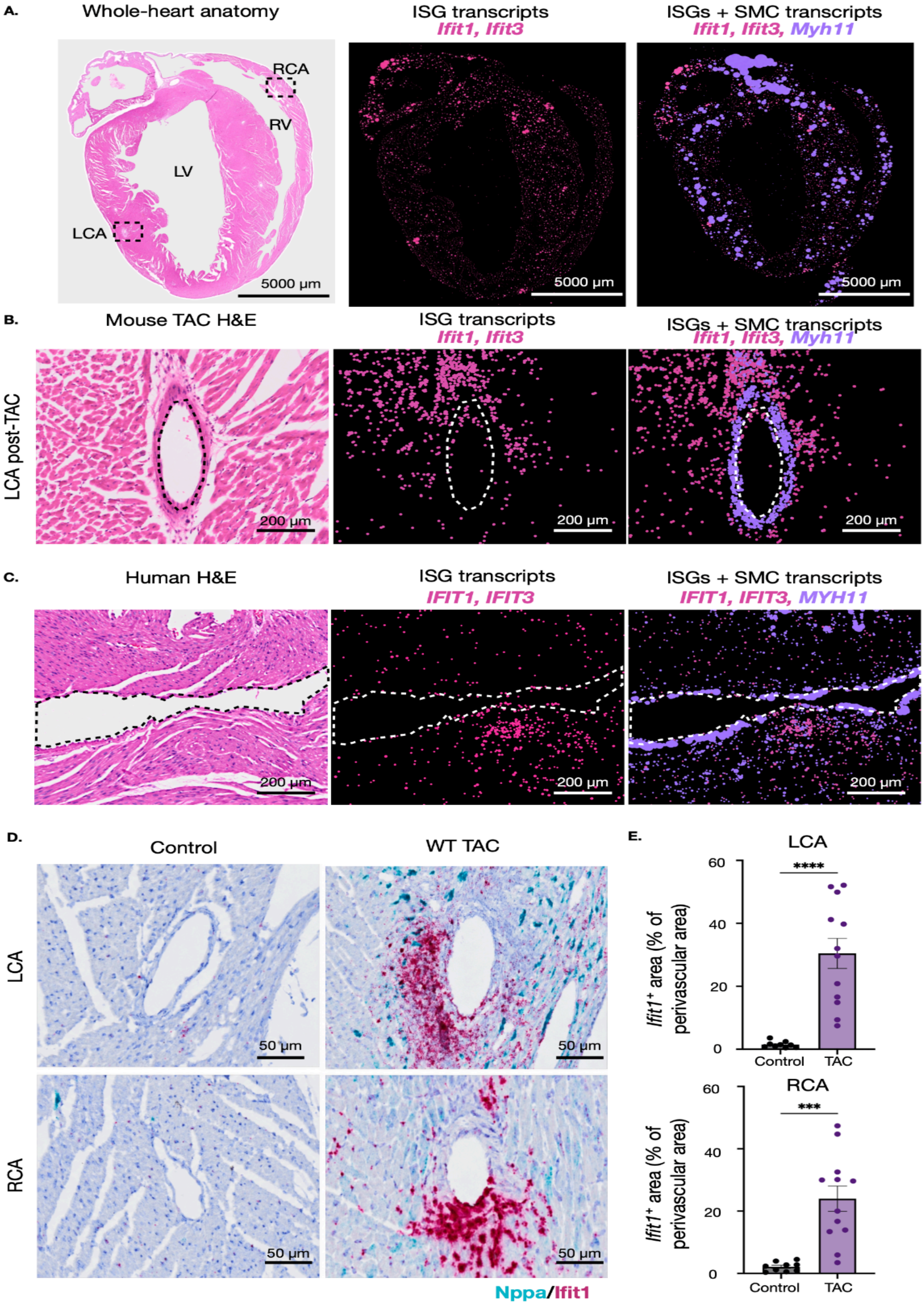

A.
Whole-heart anatomy
RCA
RV
LV
LCA
5000 µm
ISG transcripts
Ifit1, Ifit3
5000 µm
ISGs + SMC transcripts
Ifit1, Ifit3, Myh11
5000 µm
B.
Mouse TAC H&E
LCA post-TAC
200 µm
ISG transcripts
Ifit1, Ifit3
200 µm
ISGs + SMC transcripts
Ifit1, Ifit3, Myh11
200 µm
C.
Human H&E
200 µm
ISG transcripts
IFIT1, IFIT3
200 µm
ISGs + SMC transcripts
IFIT1, IFIT3, MYH11
200 µm
D.
Control
WT TAC
LCA
RCA
50 µm
Nppa/Ifit1
E.
LCA
****
Ifit1+ area (% of perivascular area)
60
40
20
0
Control
TAC
RCA
***

**Figure 3. Spatially localized ISG expression in mouse and human cardiac vessels**

(**A**) Representative whole-heart mouse Xenium spatial transcriptomic images showing H&E staining (left), Ifit1 and Ifit3 transcripts (magenta; middle), and their distribution relative to the vascular smooth muscle-associated transcript *Myh11* (purple; right). Dashed boxes in the H&E image indicate the left coronary artery (LCA) and right coronary artery (RCA) regions. LV, left ventricle; RV, right ventricle. Scale bars, 5,000 μm. (**B**) Representative mouse images of a coronary artery 7 days post TAC. H&E staining (left) illustrates vessel morphology, while spatial transcript (Xenium) maps show the distribution of ISG-associated transcripts (*Ifit1* and *Ifit3*; magenta) alone (middle) or together with the vascular smooth muscle-associated transcript *Myh11* (purple; right). Dashed outlines indicate the vessel lumen. Scale bars, 200 μm. (**C**) Representative human Xenium spatial transcriptomic images showing H&E staining (left), ISG-associated transcripts *IFIT1* and *IFIT3* (magenta; middle), and their distribution relative to the smooth muscle-associated transcript *MYH11* (purple; right). Dashed outlines indicate the vessel lumen. Scale bars, 200 μm. (**D**) Representative RNAscope images showing *Ifit1* expression in the left coronary artery (LCA; top) and right coronary artery (RCA; bottom) territories of control and wild-type TAC mice. Scale bars, 50 μm. (**E**) Quantification of Ifit1-positive area normalized to total perivascular area in the LCA (top) and RCA (bottom) of control and TAC mice. Data are presented as mean ± SEM, with individual points representing biological replicates. Statistical significance is indicated as ***P < 0.001 and ****P < 0.0001. Exact sample sizes and statistical details are provided in the corresponding Methods section.

Figure 4.

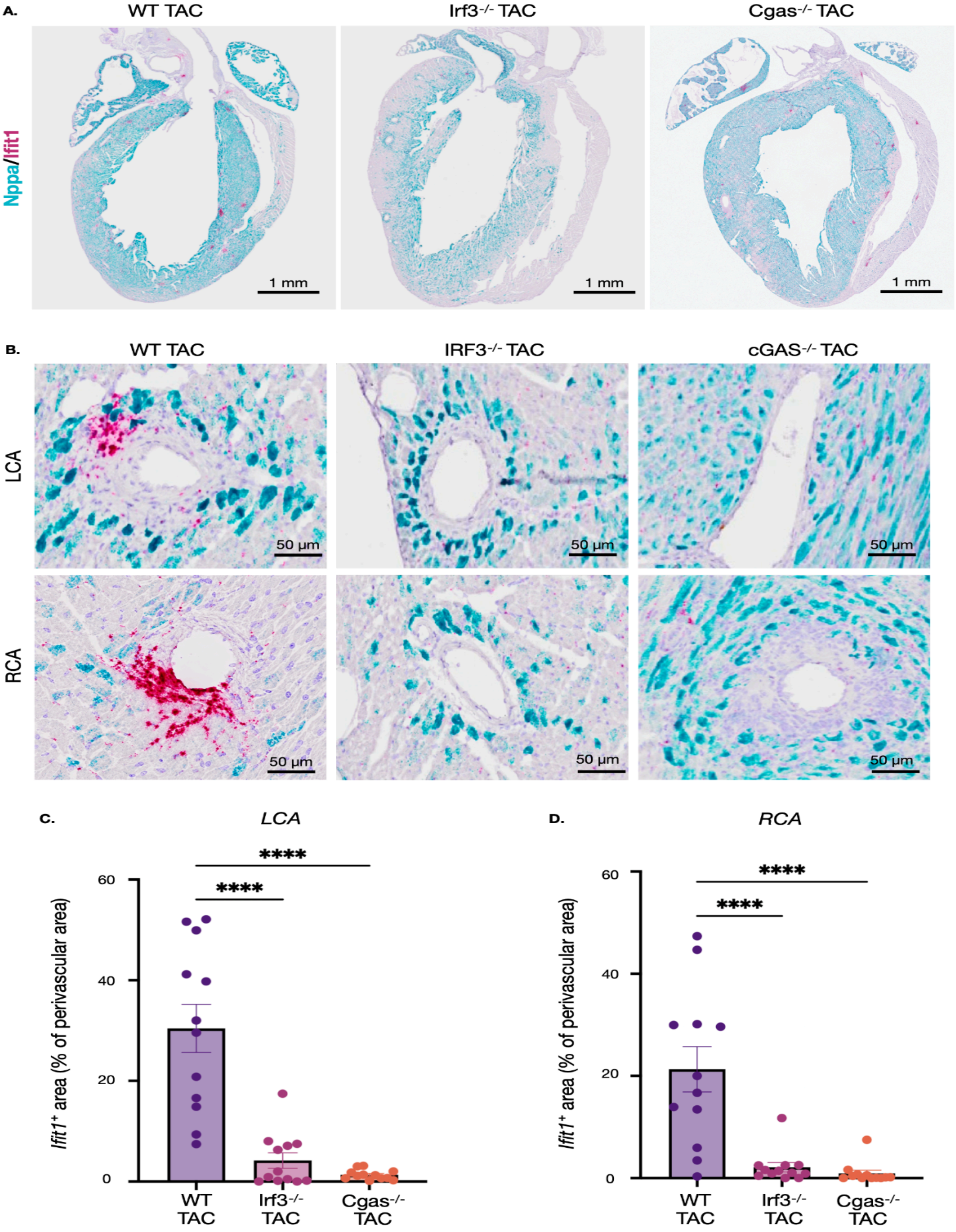
A.
WT TAC
Irf3-/- TAC
Cgas-/- TAC
Nppa/Ifit1
1 mm
1 mm
1 mm
B.
WT TAC
IRF3-/- TAC
cGAS-/- TAC
LCA
RCA
50 µm
C.
LCA
****
****
Ifit1+ area (% of perivascular area)
60
40
20
0
WT TAC
Irf3-/- TAC
Cgas-/- TAC
D.
RCA
****
****
Ifit1+ area (% of perivascular area)
60
40
20
0
WT TAC
Irf3-/- TAC
Cgas-/- TAC

**Figure 4. cGAS and IRF3 are required for robust coronary perivascular Ifit1 induction following mechanical stress.**

(**A**) Representative whole-heart RNAScope images showing *Nppa* (cyan) and *Ifit1* (magenta) expression in WT, Irf3 knockout (*Irf3*$^{-}$/$^{-}$), and Cgas knockout (*Cgas*$^{-}$/$^{-}$) hearts following TAC. *Nppa* expression is observed across all three genotypes, consistent with a cardiac stress response. Scale bars, 1 mm. (**B**) Representative high-magnification RNAscope images showing *Nppa* (cyan) and *Ifit1* (magenta) expression surrounding coronary vessels in the left coronary artery (LCA) and right coronary artery (RCA) territories in WT, *Irf3*$^{-}$/$^{-}$, and *Cgas*$^{-}$/$^{-}$ TAC hearts. Scale bars, 50 µm. (**C–D**) Quantification of Ifit1-positive area surrounding the LCA (**C**) and RCA (**D**), expressed as a percentage of the perivascular area. Perivascular Ifit1 expression is markedly reduced in Irf3$^{-/-}$ and Cgas$^{-/-}$ mice compared with WT mice following TAC. Data are presented as mean ± SEM, with individual data points representing individual vessels. Statistical analyses were performed using one-way ANOVA, ****P < 0.0001. Exact sample sizes and statistical details are provided in the corresponding Methods section.

**Figure 5.**

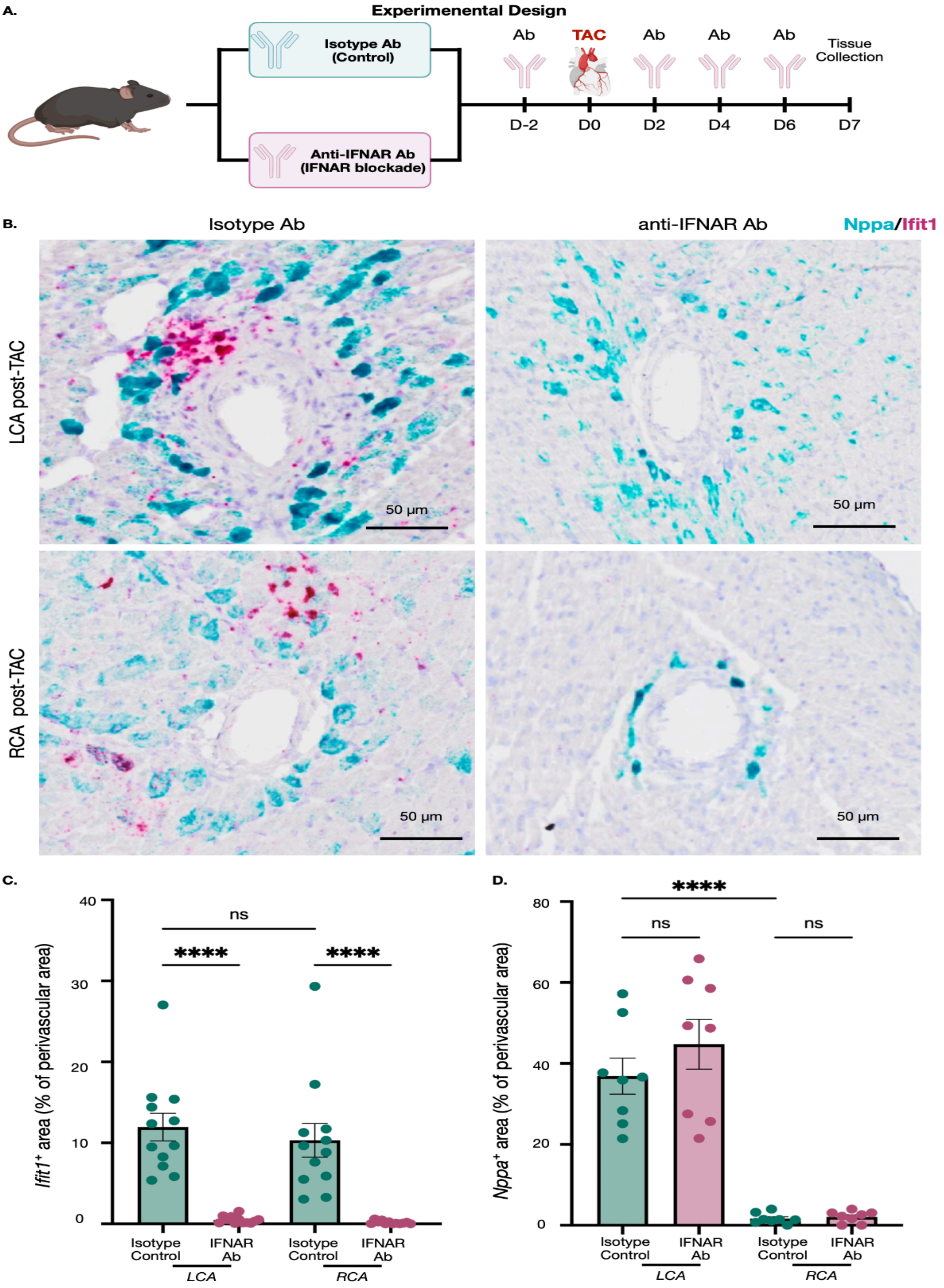
A.
Experimenental Design
Isotype Ab
(Control)
Anti-IFNAR Ab
(IFNAR blockade)
Ab
TAC
Ab
Ab
Ab
Tissue
Collection
D-2
D0
D2
D4
D6
D7
B.
Isotype Ab
anti-IFNAR Ab
Nppa/Ifit1
LCA post-TAC
RCA post-TAC
50 µm
50 µm
50 µm
50 µm
C.
Ifit1+ area (% of perivascular area)
40
30
20
10
0
ns
****
****
Isotype Control
IFNAR Ab
Isotype Control
IFNAR Ab
LCA
RCA
D.
Nppa+ area (% of perivascular area)
80
60
40
20
0
****
ns
ns
Isotype Control
IFNAR Ab
Isotype Control
IFNAR Ab
LCA
RCA

**Figure 5. IFNAR signaling is required for coronary perivascular ISG induction following mechanical stress.**

(**A**) Schematic of experimental design. Mice received isotype control or anti-IFNAR antibody (500 μg, intraperitoneally) on days −2, +2, +4, and +6 relative to transverse aortic constriction (TAC), with tissue collected on day 7. (**B**) Representative RNAscope images showing *Nppa* (cyan) and *Ifit1* (magenta) expression in the left coronary artery (LCA) and right coronary artery (RCA) territories of TAC mice treated with isotype control or anti-IFNAR antibody. Scale bars, 50 μm. (**C**) Quantification of *Ifit1*-positive area expressed as a percentage of the perivascular area in LCA and RCA territories of isotype control or anti-IFNAR antibody treated mice. **(D)** Quantification of *Nppa*-positive area expressed as a percentage of the perivascular area in LCA and RCA territories of isotype control or anti-IFNAR antibody treated mice. Each dot represents individual vessel. Data are presented as mean ± SEM. Statistical significance was determined using one-way ANOVA. ns, not significant; ****$P < 0.0001$. Exact sample sizes and statistical details are provided in the corresponding Methods section.
